# Micro-circulating hyperdynamic blood flow as a key pathogenic factor in early sepsis

**DOI:** 10.1101/2023.06.04.543593

**Authors:** Xinghuai Feng, Wei Liu, Y.B. Sun, Y. Zeng, Bu-Wei Yu

## Abstract

**Objective:** The pathogenesis of sepsis is still unknown. Sepsis 3.0 points out that “how to define sepsis and septic shock itself is still a challenge”. This study confirmed the inevitability and universality of Hyperdynamic microcirculation in sepsis, and put forward the detoxification mechanism of Hyperdynamic blood flow and the “Xinghuai Feng-Bernoulli warm shock” mechanism, that is, the pathogenic mechanism of sepsis.

**Methods:** Sepsis models of pigs, rabbits, Mice, rats and sheep were established by intravenous injection of lipopolysaccharide (LPS) and cecal ligation and perforation (CLP), and the changes of sublingual microcirculation velocity in the same branch before and after modeling were detected. SD rat model of mild sepsis was established to verify that the acceleration of blood flow is the manifestation of immune detoxification mechanism.

**Results:** The blood flow in the same branch was accelerated after the animal sepsis model was established, which was more than doubled on average. The microcirculation blood flow accelerated before the change of cardiac output CO. Rats entered a toxic state after the rapid blood flow occurred, but they could heal themselves.

**Conclusion:** The acceleration of microcirculation blood flow in sepsis is inevitable and universal, which is the cause of high output and low resistance of sepsis, and has the functions of accelerating detoxification and immunity. However, due to Bernoulli effect, it will cause oxygen exchange disorder, which is named “Xinghuai Feng-Bernoulli warm shock”, ultimately leading to hypoxia. This is the primary pathogenic mechanism of early sepsis. Since the completion of this article in 2023, its principles have undergone multicenter, randomized, and double-blind clinical verification, achieving satisfactory results. The specifics will have to wait for the publication by the three Class A tertiary hospitals.

## Introduction

Sepsis is a group of symptoms which are caused by infection, and can induce life-threatening organ dysfunction; it is a clinical syndrome with a high fatality rate. Sepsis not only seriously threatens human health, but also brings about huge economic burden on medical health. As shown by a Meta analysis of relevant studies on incidence rate and case fatality rate of adult sepsis in 27 developed countries in the period of 1979-2015, the annual incidence rate of sepsis was 288/100,000; in the past 10 years, the annual incidence rate of sepsis was about 437/100,000, with a case fatality rate of about 17%; the annual incidence rate of severe sepsis was about 270/100,000, with a case fatality rate of about 26% (Wang Zhong et al, 2020) [1]. Notably, Chinese relevant studies showed a higher case fatality rate of sepsis than that in developed countries (Wang Zhong et al, 2020) [1]. Although human research on sepsis has continuously expanded, and has entered the level of molecular biology and genomics, the pathogenesis for sepsis is yet unclear. Just as specified in the “International Consensus on Sepsis (3.0)” published in *Journal of the American Medical Association* (JAMA, February 2016), it is currently still a challenge to define sepsis and septic shock; first and foremost, sepsis is still a broad term applied to describe a process that has yet to be completely understood (Mervyn Singer et al, 2016) [2].

Under our sublingual micro-circulating imaging of sepsis patients, hyperdynamic blood flow was found, with a flow rate of >1000 μm/s, and generally 1500-3600 μm/s. These results are consistent with the results for sublingual micro-circulating imaging of 2 severe sepsis cases which were issued in the International Round-table Conference Report (2007); within the visual field in these two cases, the highest blood flow rate was 2622 μm/s and 1529 μm/s respectively (De Backer et al, 2007) [4]. See Attachment I.

By identifying the hyperdynamic blood flow in human microvessels, the initial threshold value of hyperdynamic blood flow is defined by us as 1.5 times the normal microvessel blood flow rate; the reasoning for this is given in Attachment II.

As proven by the study of (Verdant C et al, 2005) [32] and (Liu Wei et al, 2017) [33], under the condition of normal abdominal pressure, sublingual micro-circulating is highly related to visceral micro-circulating in animals. Therefore, in our study, the phenomenon of blood flow acceleration was mainly measured in sublingual micro-circulating of infection or sepsis in animals (such as pigs, sheep, rabbits and rats); possible reasons for said phenomenon and the detoxifying effect were also analyzed. As found by results of our study, the sublingual micro-circulating hyperdynamic blood flow in early sepsis found in clinical practice is a necessary and universal phenomenon when the sepsis occurs in mammals (including: human body); although it is a detoxifying process of body immune defense reaction, it causes Xinghuai Feng-Bernoulli warm shock. These findings are of great innovative significance for clarifying the pathogenesis and clinical diagnosis of sepsis.

### Take-home message

The acceleration of blood flow in the microcirculation during sepsis is an unavoidable and widespread phenomenon. It is responsible for the high output and low resistance observed in sepsis and plays a role in accelerating detoxification and boosting immunity. However, due to the Bernoulli effect, it can lead to oxygen exchange disorder, which is referred to as “Xinghuai Feng-

Bernoulli warm shock”, ultimately resulting in hypoxia. This represents the main pathogenic mechanism of early-stage sepsis.

## Methods

### Experimental animals

a: Guangxi Bama minipigs: 1 pig, provided by Laboratory for Animal Genetics and Breeding of Guangxi University, License No. SCXK(Gui)2018-0003, male, weight 27 kg.

b: Experimental sheep: 1 sheep, normal grade, male, age 18 months old, weight 26 kg, License No. SYXK(Lu)2021-0035, provided by HILE Biological Products (Shandong) Co., Ltd.

C: Japanese white rabbits: 22 rabbits, normal grade, provided by Pizhou Dongfang Breeding Co., Ltd., License No. SCXK(Su)2014-0005, male, weight 2.0-2.5 kg. These animals were randomly divided into a control group (6 rabbits) and LPS group (16 rabbits).

d. SD rats: Provided by Changzhou Cavens Experimental Animals Co., Ltd., License No. SCXK(Xu)2016-0010, 3 male rats; weight: 434 g (No.1 rat), 410 g (No.2 rat), 455 g (No.3 rat). All animal experiments conformed to relevant stipulations of Chinese governments on welfare and ethics of experimental animals(GB/T 35892-2018).

### Animal experiments of Class I

#### A: Pig LPS model

In order to observe the phenomenon of micro-circulating blood flow acceleration and its correlation with sepsis progression, LPS (Sigma Co.) was injected intravenously. In Bama minipigs, 3% Pentobarbital was continuously injected intravenously at 8-10 mL/h to maintain anesthesia; and LPS 5 μg/kg was intravenously injected. Before and after LPS injection, the variation of blood flow rate in the same branch was observed and recorded in video; a comparison analysis was made for the variation of blood flow rate. After LPS injection, observation of sublingual micro-circulating was immediately begun.

#### B: Sheep CLP sepsis model

In order to explore the time progression for the phenomenon of high output and low resistance in macro-hemodynamics of sepsis and for the phenomenon of sublingual micro-circulating hyperdynamic blood flow in its micro-hemodynamics, cecum ligation and puncture (CLP) was performed on sheep. The experimental sheep were weighed. Into the muscle of hind legs, a Midazolam Injection 0.4 mg/kg and Sufentanil Citrate Injection 0.007 mg/kg was injected to sedate the subjects. After the animals fell asleep, they were kept in a supine position through protective constraints. After tracheotomy of the anterior cervical region, a #7.0 tracheal catheter was implanted. A ventilator was connected for mechanical ventilation (SC-3 invasive ventilator, Nanjing Puao Medical Equipment Co. Ltd., China). Through accessing the internal jugular vein, a double-lumen central vein catheter was implanted. Through this internal jugular vein, a Propofol Injection of 15 mg/kg/h and a Rocuronium Bromide for Injection 0.2 mg/kg were given for maintenance anesthesia. Parameters for the ventilator were adjusted as follows: ventilation mode of volume control; tidal volume 300 mL; positive end-expiratory pressure (PEEP) 5 cmH2O; fraction of oxygen inspiration (FiO2) 1; inspiratory to expiratory ratio (I/E) 1:2. An ECG monitor (Dash 2000, GE, GuoXieZhuZhun 20058730009, China) was connected for measurement of heart rate and non-invasive blood pressure. A sensor (FloTrac MHD8 cardiac output and Pressure Monitoring Sensor, Edwards Lifesciences LLC, Canada) was connected with the internal jugular vein for measurement of the central venous pressure (CVP). A Hemodynamic monitor (Vigileo MHM1E, Edwards Lifesciences, GuoXieZhuZhun 20063211962, USA) was connected with common carotid for measurement of cardiac output and cardiac index. Then, a septic shock model was established through CLP surgery. When mean arterial pressure dropped to 70% of normal blood pressure, the establishment of the septic shock model was considered successful. After successful modeling, observation of the sublingual micro-circulating was begun. At observation, all secretes were first cleared off of sublingual mucosa. Then, the LH-SDF-2 sidestream dark-field (SDF) vital microscope with disposable transparent protective sheath in the front of the probe was gently adhered to the sublingual lateral side (Note: Avoid an obvious compression on the mucosa, so as not to influence the vascular engorgement. Adjust the focal length for obtaining of clear images).

Through the above vital microscope, software and Multi-angle adjustable bracket observations of whether the images of Blood flow in the same blood vessel acceleration appeared in sublingual micro-circulating of sheep before and after the modeling were made.

#### C: Rabbit LPS model

Japanese white rabbits were raised for one week in a desensitized environment. Before the surgery, they were made to fast for 24 hours. Into the vein at the margin of the ear, 20% Urethane 4-6 mL/kg was injected for anesthesia. After the tracheotomy, a tracheal catheter was implanted. A 24G polyethylene catheter was implanted into the common carotid. Through a pressure energy converter, a biological signal collection system (BL-420S, Chengdu Taimeng Instrument Co., Ltd.) for thalline alliance in Chengdu) was connected for continuous monitoring of invasive arterial blood pressure.

In the LPS group, LPS 2 mg/kg (Sigma Co.) was injected into the vein at the margin of the ear; in the control group, LPS was not injected, but only normal saline of equal amount was injected. When mean arterial pressure dropped to 70% of blood pressure before LPS injection, the occurrence of sepsis was considered to have happened. By localizing through the LH-SDF-2 SDF vital microscope, it was located on a microvessel larger than 20μm. After the LPS injection, the observation was started. At the time points of 0min, 5min, 10min, 15min, 20min, 25min, and 30min, videos were collected; the animals were kept immobile until the completion of video recording.

#### D: Mice LPS model

**Variety/strain: Male BALB/c mice; Grade: SPF grade; Supplier: Sichuan University Experimental**

### Animal Center

**Heart; Production license number: SCXK (Chuan) 2018-026; Gender and quantity of purchased animals: 48, half male and half female, weight: 20g**.

#### 1.1.1 Animal feeding

**Breeding location: Experimental Animal Center of Sichuan University; Animal Use License Number: SYXK (Chuan) 2018-185; Environmental level: Barrier system; Temperature: 20-25** □**; Relative humidity: 40∼70%; Ventilation rate: 10-15 times/hour; Lighting time: alternate lighting every 12h/12h. Mouse cage: B3 type transparent mouse cage (L × W × H: 37 mm × 28 mm × 11 mm), produced by Suzhou Fengshi Experimental Animal Equipment Co**., **Ltd**., **with 5 animals per cage. Mouse bedding: The bedding is corn cob, provided by Beijing Ke’ao Xieli Feed Co**., **Ltd**., **and is sterilized by high temperature and high pressure steam before being used for animal use. Update frequency of mouse cages: 2 times a week. Cleaning and disinfection: Clean up after daily experimental operations and feeding work, and then disinfect in the animal management room. Large/mouse full price pellet feed, produced by Beijing Ke’ao Xieli Feed Co**., **Ltd**., **license number: SCXK (Jing) 2019-0003, in compliance with B14924.3-2001 standard. Feeding method: Free ingestion. During the experiment, rats were given high-pressure steam sterilized tap water, which was freely ingested from animal drinking bottles**.

### 1.2 Model Preparation and Processing

#### 1.2.1 Construction and experimental grouping of early sepsis experimental model

**SPF grade BALB/c male mice (weighing approximately 20g) were randomly divided into: A. Control group (physiological saline group); B. 0.5 mg/kg LPS group; C. 2.5 mg/kg LPS group; D. 5mg/kg LPS group. By injecting different concentrations of lipopolysaccharide (LPS) into the tail vein of mice, the microcirculation of the right ear auricle was dynamically monitored, with a focus on observing the trend of microcirculation blood flow changes, detecting the time window of hyperdynamic blood flow generation in microcirculation, clarifying the occurrence pattern of hyperdynamic blood flow in microcirculation, and clarifying the dependence of hyperdynamic blood flow in microcirculation on LPS concentration**.

### 1.3 Mouse Ear Microcirculation Imaging Collection

**Firstly, apply an appropriate amount of cedarwood oil on the surface of the experimental platform. After anesthesia with isoflurane, the inner side of the right ear of the mice is tightly adhered to the experimental site**.

**On the surface of the table, soak a cotton swab in an appropriate amount of cedarwood oil, and then use the cotton swab to smooth the earlobes along the hair direction. Using a microvascular collateral flow dark field imaging observation system to detect changes in microvascular blood flow, adjust the focal length, and collect clear and stable image data. After data collection is completed, the blood flow velocity is measured using the Microvascular Flow Dark Field Imaging Observation System (XUZHOU LIHUA ELECTRONIC TECHNOLOGY DEVELOPMENT CO**., **LIMITED, LH-SDF-2) and recorded for analysis**.

### 5. Measurement of microcirculation blood flow velocity in mouse auricle

**Measure the microcirculation blood flow velocity in the mouse auricle using the image analysis tool of the Microvascular Flow Dark Field Imaging Observation System (LH-SDF-2). Open the image to be tested, mark the blood vessel to be tested, select the flow velocity analysis tool and use the comparison method to measure the blood flow velocity of the vessel. When the flow velocity is too fast, use the flow velocity tracking measurement tool to measure the blood flow velocity of the vessel, record the diameter and flow velocity of the measured vessel, and continuously measure the blood flow velocity of the same branch of the vessel multiple times over time. The blood flow velocity of the blood vessels is divided into stages T every 3 minutes, with T0 as the LPS injection time point. The final measurement result is represented by the mean blood flow velocity within stage T**.

### Animal experiments of Class II: detoxifying mechanism test

#### Rat LPS model

In order to assess the influence of hyperdynamic blood flow on the detoxifying ability of animals, LPS of different doses was injected into rats. After weighing the 3 rats, LPS 0.5 mg/kg (L8880, Solarbio) was injected into their caudal vein; with the dose being based on weight: 434 μL for No. 1 rat, 410 μL for No.2 rat, and 555 μL for No.3 rat (i.e. by an increase of 100 μL over the weight-based dose).

#### Measurement of sublingual micro-circulating

Through a LH-SDF-2 hand-held vital microscope independently developed by Xuzhou Lihua Electronic Technology Development Co., Ltd., sublingual micro-circulating images were collected; the video recording lasted about 20 seconds each time. At video recording, the instrument was placed and fixed into a Multi-angle adjustable bracket in cohesion with the vital microscope; then, the probe of the instrument was gently placed into the sublingual mucosa; gas bubbles under the lens were eliminated through normal saline gauze. The pressure was adjusted until larger microvessel blood flow and more obvious images of blood cell continuous flow were clearly found, so as to eliminate artifacts. Finally, the same microvessel was localized, which was then kept immobile until the completion of testing.

#### Analysis of micro-circulating blood flow rate

In micro-circulating images, clear erythrocytes or leukocytes in sublingual micro-circulating branches were traced. Through the independently-developed Chinese Advanced Microvessel Analysis & Comparison System software (Version 1.0, Xuzhou Lihua Electronic Technology Development Co., Ltd.), the passing distance of these blood cells within a certain time was analyzed to obtain the micro-circulating mean blood flow rate. In our study, the hyperdynamic blood flow rate was not measured using the current universal space-time method, the reason for this is given in “Discussion”. [10]

## Statistical analysis

The measurement data were analyzed using SPSS 25.0 software and presented as mean ± standard deviation (mean ± SD). Differences among multiple groups were assessed by one-way analysis of variance (ANOVA). A p-value < 0.05 was considered statistically significant.

## Results

(1) Animal experiments of Class I: The phenomenon of micro-circulating blood flow acceleration was reproduced; it was proven as necessary. It was determined that such a phenomenon occurred in the early period of infection.

After LPS injection into pig sublingual micro-circulating, the vessel of the same branches sublingual micro-circulating blood flow was gradually accelerated over 3-9 minutes. sublingual micro-circulating blood flow was gradually accelerated over 3-9 minutes. Concrete measurement and comparison before and after LPS injections are shown in Figure 1.

**Figure 1:**
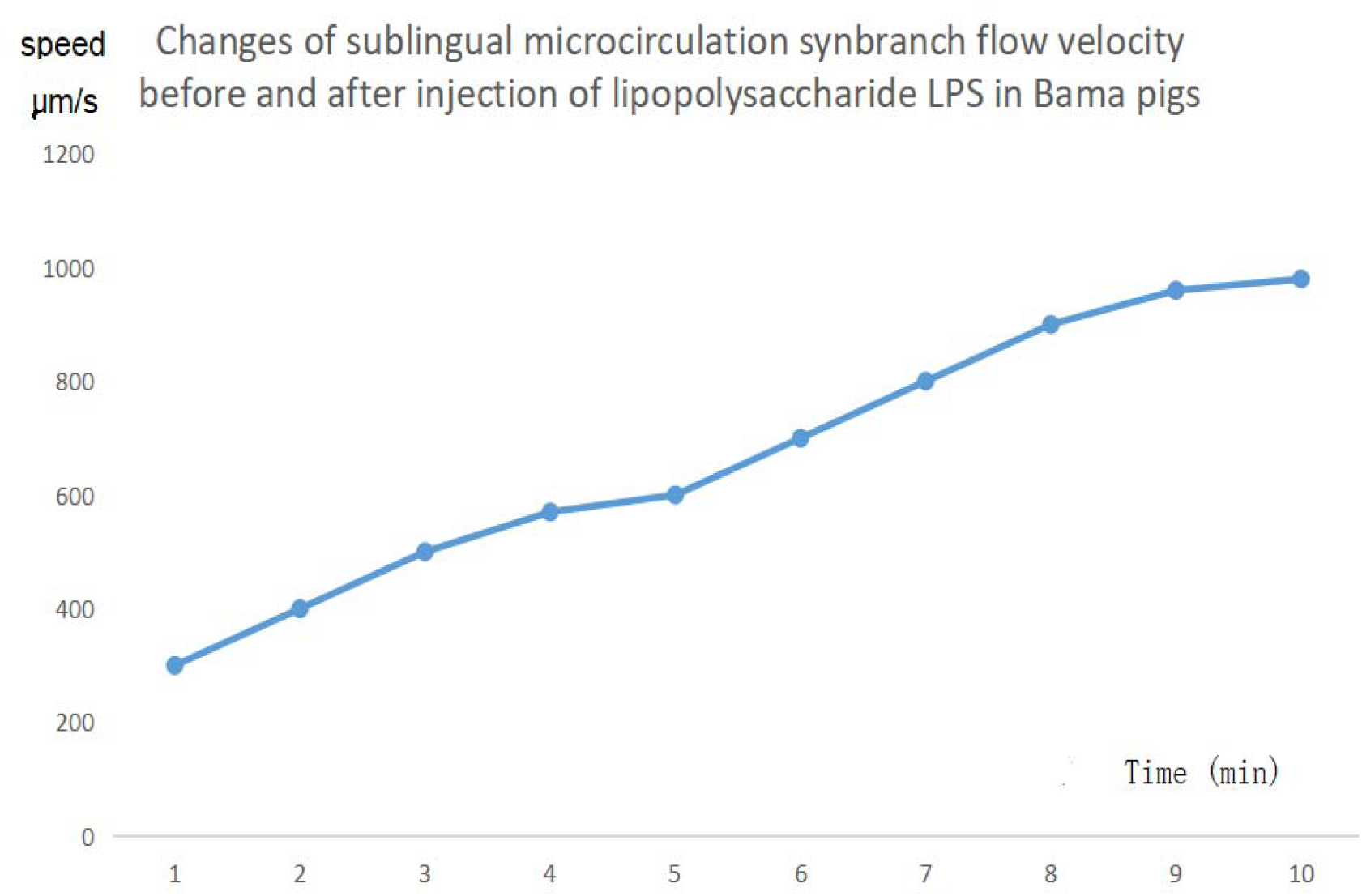
Changes of sublingual microcirculation synbranch flow velocity before and after injection of lipopolysaccharide LPS in Bama pigs. After LPS 5 μg/kg (Sigma Co.) was intravenously injected into Bama pigs, the blood flow velocity of the same branch of blood vessel gradually rose from 302 μm/s before the injection (Time 1) to 560 μm/s (Time 5, 5min), peaking at 958 μm/s (Time 9, 9min). Y-axis: Flow rate; X-axis: Time.

(2) The micro-circulating hyperdynamic blood flow in micro-hemodynamics was related to the macro-hemodynamic indices of cardiac output and cardiac index, and the phenomenon of blood flow acceleration was reproduced again; proving the necessity for its occurrence.

After CLP infection of sheep, the time variation was monitored for sublingual micro-circulating Blood flow velocity of the same branch of blood vessels, heart rate, cardiac output, cardiac index, mean arterial pressure and central venous pressure under continuous branch sublingual micro-circulating blood flow. Concrete detail is shown in Figure 2.

**Figure 2:**
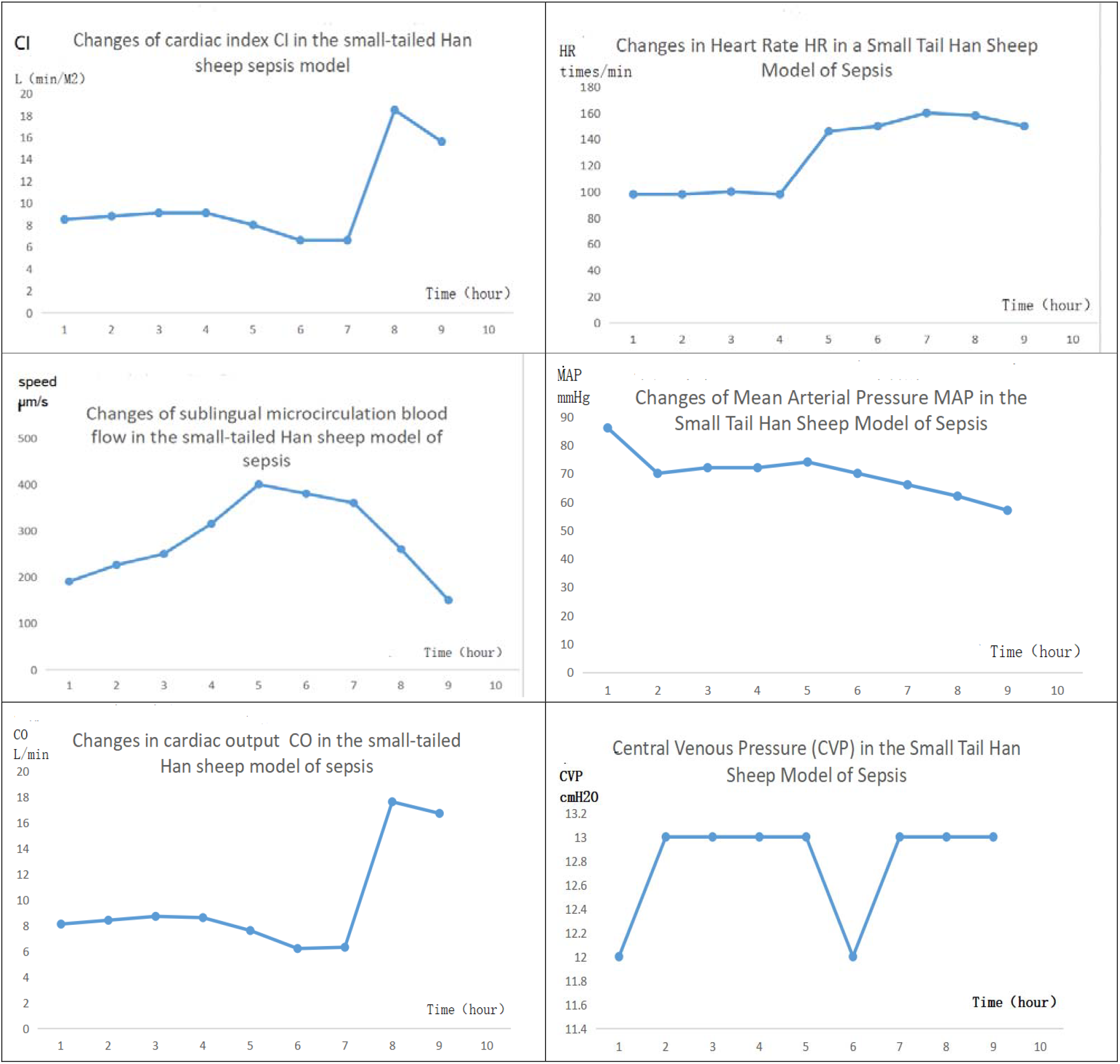
Time variation of the speed of sublingual micro-circulating blood flow, heart rate, cardiac output, cardiac index, mean arterial pressure, and central venous pressure during the establishing course of experimental sheep sepsis model.

Surgery began at 10:00 on day 1. At 11:40 of the next day, the model was established successfully. At 4:00 of the next day, measurement began. At the completion of surgery one day 1, the speed of sublingual micro-circulating blood flow, heart rate, cardiac output, cardiac index and mean arterial pressure (MAP) was measured; MAP was not significantly different from that of 4:00 the next day. Therefore, from 4:00 of the next day, the corresponding variation at the same time point was compared for the speed of sublingual micro-circulating blood flow, heart rate, cardiac output, and cardiac index. As shown by Figure 2, sublingual micro-circulating blood flow reached 190 μm/s at 4:00 of the next day (Time 1), 250 μm/s at 6:00 (Time 3), maximum 400 μm/s at 8:00 (Time 5), 350 μm/s at 10:00 (Time 6), and 150μm/s at 11:40 (Time 8.5) when the criteria for successful modeling was met. From Time 1 to Time 5, cardiac output was not increased synchronously with micro-circulating blood flow; it started to rise from Time 7 to Time 8. The variation trend of the cardiac index was similar to that of the cardiac output. The variation of the cardiac output and the cardiac index lagged behind that of the micro-circulating blood flow, but the variation trend of the flow rate was the same as the increase trend of the cardiac output and cardiac index.

(3) Prove the inevitability of accelerated blood flow in early sepsis from a statistical perspective 22 Japanese white rabbits were divided into the endotoxin group (16 rabbits) and control group (6 rabbits). In the endotoxin group, the variation of sublingual micro-circulating blood flow rate was observed after LPS injection; in the control group (without LPS injection), the variation of sublingual micro-circulating blood flow rate was observed; results are shown in Figure 3. In the endotoxin group, the phenomenon of blood flow acceleration lasts >20 minutes; at 25min, the blood flow rate was not obviously different between the two groups; after the occurrence of shock, the blood flow rate in the endotoxin group showed an obvious decreased than that of the control group. P<0.05 indicates that the difference was statistically significant. All values were expressed with mean value ± standard deviation.

**Figure 3:**
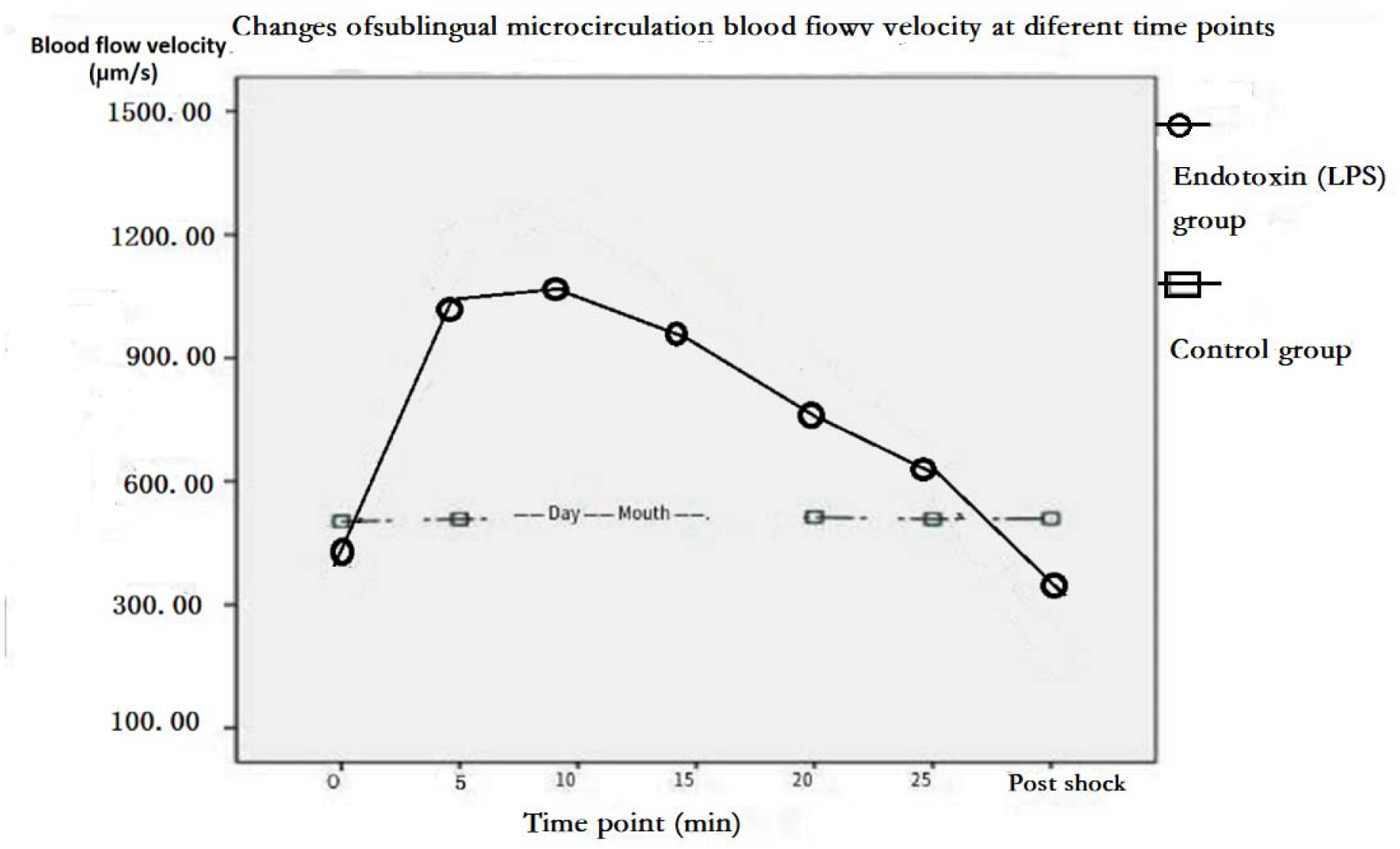
Comparison of the variation of sublingual micro-circulating blood flow rate between two groups. As more visually shown by such comparison, at 5-10 minutes after the injection, the blood flow rate in the endotoxin group was increased by >1 fold compared to the control group; in the control group, the flow rate was steady and did not increase at all.

**(4) The effect of different doses of LPS on microcirculation blood flow velocity in early sepsis mice**

**Low (0.5 mg/kg), medium (2.5 mg/kg), and high (5.0 mg/kg) doses of LPS were injected into the tail vein of BALB/c mice. The microvascular bypass dark field imaging system (LH-SDF-2) was used to monitor the changes in microcirculation blood flow in the right ear auricle of BALB/c mice. The measurement time was measured at 3-minute intervals T, with T0 as the LPS injection point. The comprehensive results are shown in Figure 1. Similarly, in the T1 stage, the blood flow velocity of the low, medium, and high-dose LPS groups significantly increased compared to the saline group, and there was no significant difference between the groups, indicating that the low, medium, and high-dose LPS groups produced hyperdynamic blood flow at the same level in the T1 stage. However, the overall changes in blood flow velocity during the T0-T9 stages were different among the low, medium, and high dose LPS groups. The blood flow velocity of the low-dose LPS group showed an increasing trend at T0-T1, a decreasing trend at T2-T9, and returned to normal levels after T9 (30 minutes). The blood flow velocity of the medium dose LPS group showed an upward trend during the T0-T9 stage, while T2-T4 maintained hyperdynamic blood flow in T1. The microcirculation blood flow velocity further increased during T5-T8, and the changes in microcirculation blood flow velocity after T9 need to be further clarified. The blood flow velocity of the high-dose LPS group showed an upward trend in the T0-T2 stage, a downward trend in the T2-T9 stage, a significant decrease in blood flow velocity compared to the T3 stage and lower than that of the saline group in the T4 stage, and continued to decrease until the T7 stage, approaching stagnation**.

**Figure 1.**
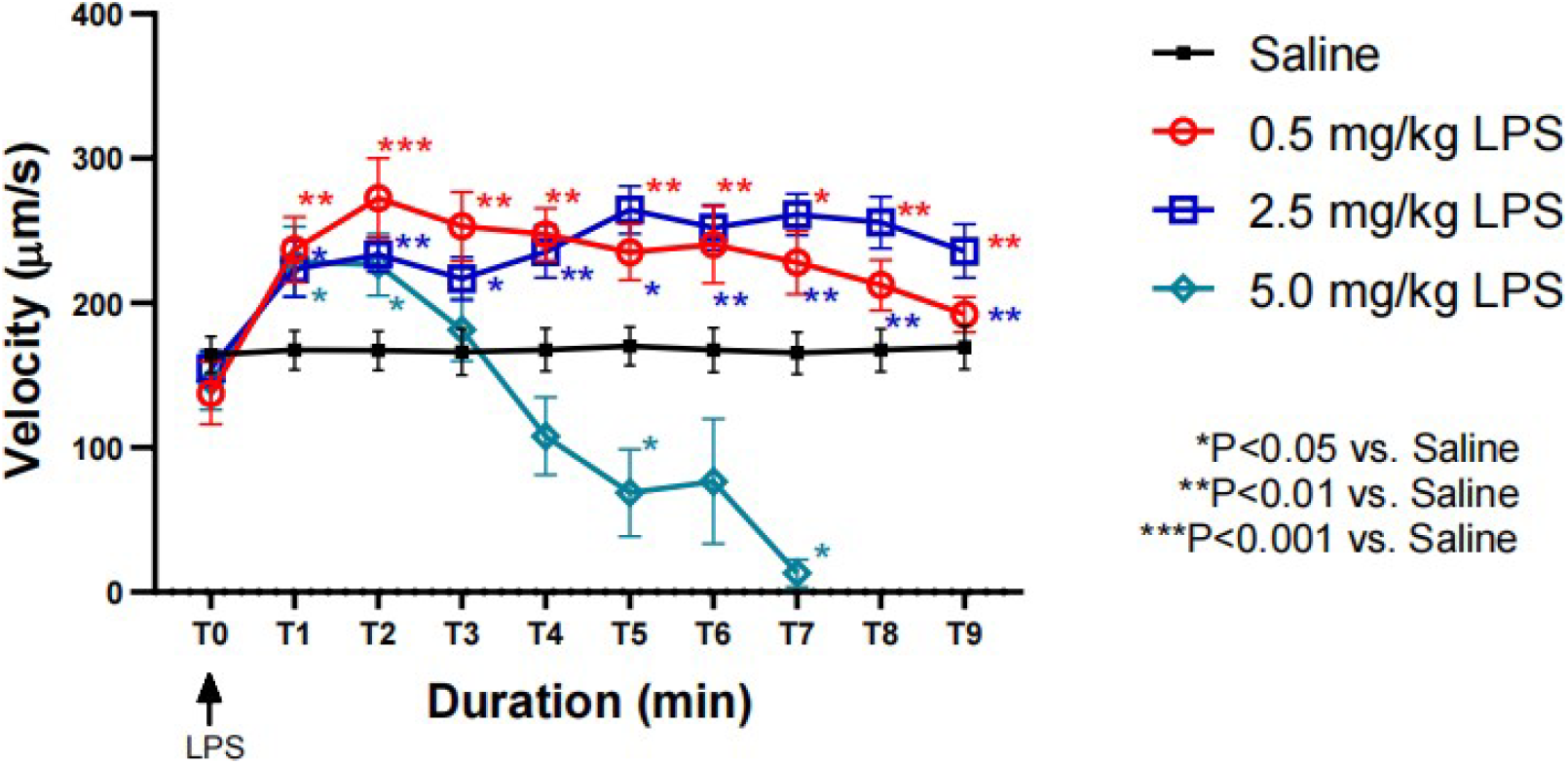
The time-dependent changes in microcirculatory blood flow velocity induced by low (0.5 mg/kg), medium (2.5 mg/kg), and high (5.0 mg/kg) doses of LPS in early sepsis* p <0.05 vs. Saline;** p <0.01 vs. Saline;*** p <0.001 vs. Saline.

The effects of 2 different doses of LPS on the appearance time and duration of microcirculatory hyperdynamic blood flow in early sepsis mice

By analyzing the three blood flow parameters of hyperdynamic blood flow occurrence time,hyperdynamic blood flow duration score, and maximum blood flow velocity

Further analysis was conducted on the effects of low, medium, and high doses of LPS on early blood flow changes in mice with sepsis, as shown in Figure 2. The time of hyperdynamic blood flow in the low, medium, and high dose LPS groups was mainly concentrated within 4-6 minutes, with no significant difference between the groups. There was nohyperdynamic blood flow in the saline group. The duration score of hyperdynamic blood flow in the medium dose LPS group was significantly higher than that in the low-dose and high-dose LPS groups, indicating that the duration of hyperdynamic blood flow in the medium dose LPS group was significantly higher than that in the low-dose and high-dose LPS groups. Similarly, the highest blood flow velocity in the medium dose LPS group was significantly higher than that in the low dose and high dose LPS groups, and the highest blood flow velocity in the low, medium, and high dose LPS groups was significantly higher than that in the saline group.

Table 1 Analysis results of hyperdynamic blood flow occurrence time,hyperdynamic blood flow duration score, and maximum blood flow velocity parameter.

**Table 1.** Analysis results of hyperdynamic blood flow occurrence time, hyperdynamic blood flow duration score, and maximum blood flow velocity parameter.

| Table 1. Analysis results of hyperdynamic blood flow occurrence time, hyperdynamic blood flow duration score, and maximum blood flow velocity parameter |  |  |  | Mean±SD |
| --- | --- | --- | --- | --- |
| group | Hyperdynamic blood flow occurrence time (min) | Hyperdynamic blood flow duration rating |  | Maximum blood flow velocity (μ m/s) |
| saline | / | 0 |  | 181.0±40.05 |
| 0.5 mg/ml LPS | 5.22 ± 0.45 | 1.64±1.22 |  | 253.4±70.07 |
| 2.5 mg/ml LPS | 4.81 ± 0.69 | 3.25±0.75 |  | 334.90±32.71 |
| 5.0 mg/ml LPS | 4.51 ± 0.94 | 1.15±0.99 |  | 240.50±74.32 |

**Hyperdynamic blood flow duration score: A hyperdynamic blood flow duration of 0 T is scored as 0, 1–3 T is scored as 1, 4–6 T is scored as 2, 7–9 T is scored as 3, and 9 T or more is scored as 4.**

**Figure 2.**
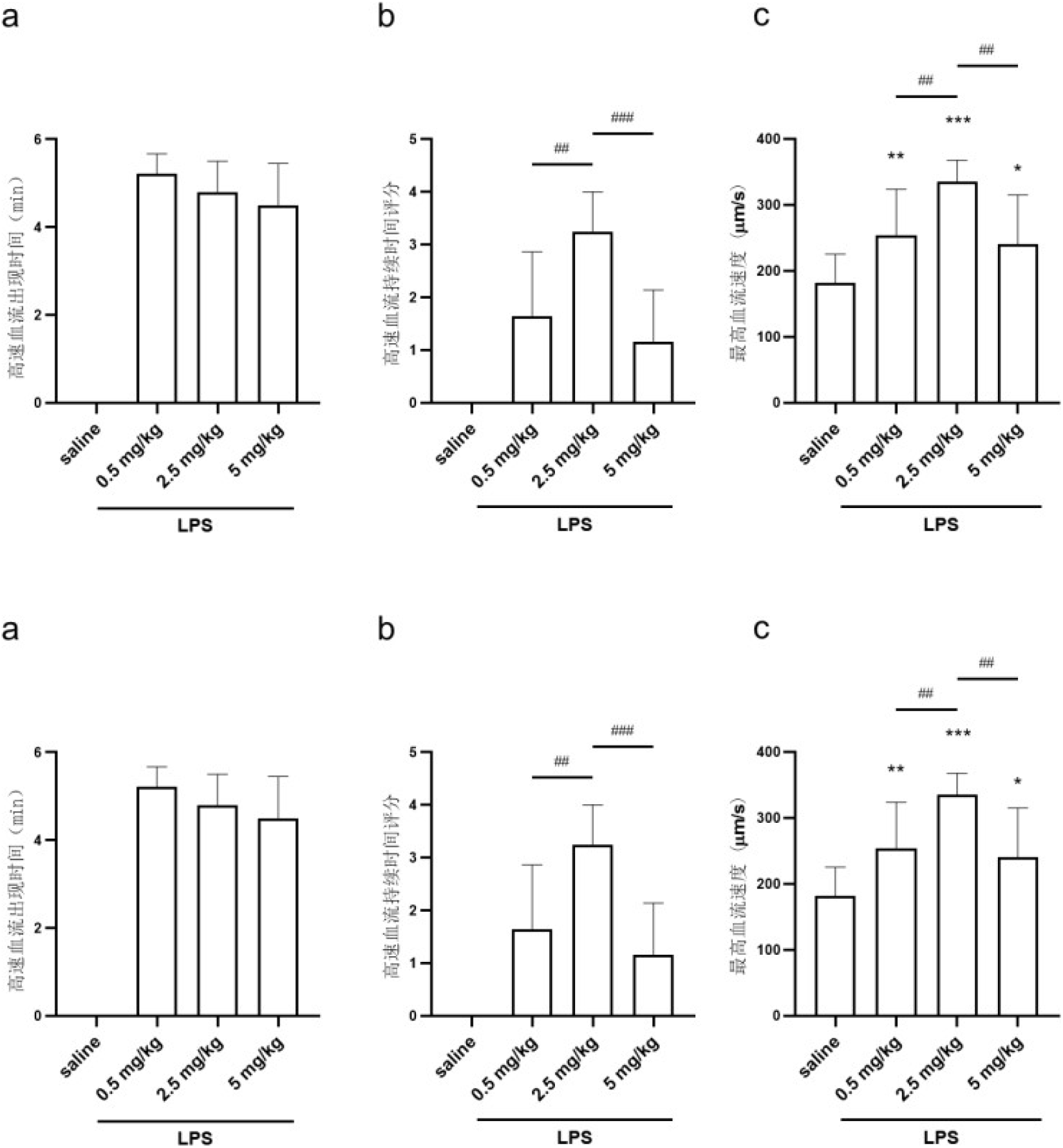
Analysis results of early microcirculatory hyperdynamic blood flow occurrence time,hyperdynamic blood flow duration score, and highest blood flow velocity parameter induced by low (0.5 mg/kg), medium (2.5 mg/kg), and high (5.0 mg/kg) doses of LPS in sepsis. a. Time analysis of hyperdynamic blood flow occurrence. b. Analysis of hyperdynamic blood flow duration score.

The generation of hyperdynamic blood flow in mouse microcirculation is inevitable after tail vein injection of different doses of LPS, but the duration and size of hyperdynamic blood flow in microcirculation are affected by LPS concentration.

Animal experiments of Class II: to validate that animal body possessed a certain detoxifying mechanism Rat No. 1; LPS injection into caudal vein: The branch sublingual micro-circulating blood flow rate reached 566 μm/s before the injection, 708 μm/s at 6 minutes after the injection, and 1113 μm/s at 9 minutes after the injection. Rat No. 2; LPS injection into caudal vein: The branch sublingual micro-circulating blood flow rate reached 130 μm/s before the injection and 340 μm/s at 8 minutes after the injection. Rat No. 3; LPS injection into caudal vein: The dose was increased by 100 μL over the weight-based dose (i.e. 455 μL + 100 μL), The branch sublingual micro-circulating blood flow rate reached 200 μm/s before injection and 460 μm/s at 11 minutes after injection.

In rats No. 1 and No. 2, after LPS injection at experimental dose under anesthetic state, sublingual micro-circulating blood flow was accelerated; after resuscitation from anesthesia, toxic symptoms occurred; the rats did not eat or drink; and the response was lagging; within 24 hours of injection, the rats were in a semi-conscious state; after 24 hours, water drinking gradually begun; after 48 hours, vigor gradually restored; after 72 hours, response to human touch was restored, and the rats were indistinguishable from normal rats. In rat No. 3, the stagnant blood flow was gradually accelerated after 8 hours; and the rat died the next day.

Therefore, when the dose of LPS exceeds the detoxifying ability of animals, the animals will die. See Table 1.

Hyperdynamic blood flow may be a manifestation of initiation of the sympathetic-immune defense function. When the toxin invades the blood beyond the reasonable limit that the blood allows, the body initiates hyperdynamic blood flow so as to rapidly transport the toxins to the liver and kidney for detoxification. Therefore, this is actually an immune defense reaction that is formed by the body over the course of evolution. This is the detoxifying mechanism for the generation of hyperdynamic blood flow.

**Table 1.** Experimental results after LPS injection at limited dose and at overdose into the caudal vein of rat No. 1 and rat No. 2 under an anesthetic state.

|  | Ctrotrol group |  | Observation group |
| --- | --- | --- | --- |
| Rat tail vein injection group | No.1 limited injection | No.2 limited injection | No.3 Overdose injection |
| Weight | 434 g | 410 g | 455 g |
| Injection quantity | 434 uL | 410 ul | 455uL+100uL |
| Blood flow velocity<br>before injection LPS into<br>the same branching<br>vessel (um/s) | 566 | 130 | 200 |
| Blood flow velocity<br>after injection LPS into<br>the same branching<br>vessel for 8-12 minutes (um/s) | 1113 | 340 | 460 |
| Post-injection state | 24-hour resumption of diet | 24-hour resumption of diet | He died 24 hours later |

## Discussion

In order to clarify the pathogenesis of sepsis, we first raised 5 questions, then answered them by combining the above experiments; then, according to experimental results, literature data, and logical judgment, detailed proof was given for these answers; finally, such information was integrated into an innovative new scientific theory for the pathogenesis of early sepsis.

1. Question 1: Is hyperdynamic blood flow in sepsis a necessary and universal phenomenon? Answer 1 and Proof 1 are provided below.

Question 1 was raised according to the found sublingual micro-circulating hyperdynamic blood flow in human sepsis: At the beginning of our study, a hyperdynamic blood flow in human sepsis was measured according to sepsis videos from relevant literature (De Backer et al, 2007) [4], (Refer to Attachment I). The question is whether hyperdynamic blood flow in sepsis is a necessary and universal phenomenon?”.

Answer 1: This phenomenon is an immune defense reaction formed by the human body over the course of evolution; the generation of hyperdynamic blood flow is the body’s detoxifying mechanism, and thus it is a necessary and universal phenomenon.

Proof 1: We first proved the necessity, and then proved the universality. Of course, a universal phenomenon is also a necessary one.

As this phenomenon is necessary and universal, it should be reproduced in animal experiments. Therefore, animal experiments of Class I and II were designed. As proven by experiments on animals of different species and different quantities, the blood flow rate of the same branching vessel after the establishment of an animal sepsis model is increased by >1 folds than before modeling. Therefore, the variation trend of sublingual micro-circulating blood flow acceleration in the early sepsis of animals is consistent with that in sepsis patients found in clinical practice; proving the judgment as necessary in Answer 1, i.e. micro-circulating blood flow acceleration (with manifestations of hyperdynamic blood flow in the human body) is a necessary phenomenon of mammals (including human beings) in early sepsis.

Proof for the universality of hyperdynamic blood flow in early sepsis: The train of thought for this proof is shown as follows: Human beings universally realize a phenomenon of high output and low resistance in the macro-hemodynamics of sepsis; through the animal experiment of Class I on sheep, literature, physiological theory, and shock theory, a correlation between the hyperdynamic blood flow and the phenomenon of high output and low resistance was verified to thus prove that the hyperdynamic blood flow is universal for the occurrence of sepsis (in fact, this is also proof for its necessity).

(1) Proof of experiment: As shown by the sequence of time axis in Figure 2 at animal experiments of Class I on sheep, micro-circulating blood flow was first accelerated, and an increase in heart rate, cardiac output, and cardiac index occurred approximately two hours after the onset of high dynamic blood flow in the microcirculation. This phenomenon indicates that the invasion of pathogenic factors was first perceived by vascular endothelial cells in peripheral micro-circulating; then, in order to avoid further injury to the human body, micro-circulating hyperdynamic blood flow is started to accelerate the detoxifying process. The micro-circulating hyperdynamic blood flow necessarily causes a large increase in venous return volume; therefore, in order to keep a dynamic balance, cardiac output is necessarily increased correspondingly. In other words, the micro-circulating blood flow acceleration is the reason for high output and low resistance in macro-hemodynamics; the micro-circulating hyperdynamic blood flow initiates the phenomenon of high output and low resistance in macro-hemodynamics. Since the micro-circulating hyperdynamic blood flow first starts slowly (i.e. the convergence and returning of vein blood to the heart gradually increases), Therefore, there is a relatively lagging process reflected in the cardiac output CO. This reveals an inherent correlation between the warm shock phenomenon of high output and low resistance in early sepsis and the phenomenon of sublingual micro-circulating hyperdynamic blood flow; it is proven that they are just different manifestations of the same pathologic factor in macro-hemodynamics and micro-hemodynamics. As concluded in the literature (Ravikant T, Walt AJ et al, 1976) [15], (Christoph Langenberg, et al, 2006) [18] and (Auio S. HERMRECK, et al, 1969) [26], hyperdynamic blood flow occurs after the establishment of sepsis model for experimental pigs, sheep and dogs; since this phenomenon was not observed together with micro-hemodynamics, their inherent correlation was not revealed. Once the correlation between the micro-circulating blood flow rate in micro-hemodynamics and the cardiac output or cardiac index in macro-hemodynamics is ascertained, the universality of micro-circulating hyperdynamic blood flow in sepsis can be further realized by utilizing the human knowledge that the high output and low resistance in macro-hemodynamics of sepsis is a universal phenomenon.

(2) Proof of literature: The high output and low resistance in the macro-hemodynamics of sepsis is a unique universal phenomenon that has been realized for sepsis by human beings. In Page 386 of the book “Prevention and Treatment of Sepsis”, Yong-Ming Yao et al specifies as follows: As shown by results of clinical observation and hemodynamic monitoring on numerous cases, septic shock patients were at a state of hyperdynamic circulation throughout most of the illness (i.e. cardiac output was normal or higher than normal value; total peripheral resistance was decreased); some patients were also at a hypodynamic in the latter period (i.e. low output and high resistance) (Yaoyong Ming et al, 2018) [3]. In Page 125 of the physiological textbook (9^th^ edition), the following contents are mentioned in the part “blood circulation-micro-circulating”: At infective or toxic shock, arteriovenous shunt and thoroughfare channel are opened in large number; although the patients are in a state of shock, as the skin is warmer (i.e. warm shock is considered at this time); since a large amount of micro arterial blood enters through anastomotic branch into the micro vein and does not make material interchange with histiocytes, tissue anoxia can be aggravated to exacerbate the illness state (Zhuda Nian et al, 2018) [11]. Therefore, at infective or toxic shock, warm shock is a universal phenomenon. One piece of research (Liuda Wei et al, 2013) [5] states as follows: As a special type of shock, the infective shock usually has a sign of high cardiac output and low peripheral vascular resistance in macro-hemodynamics. In much of the foreign literature, the phenomenon of high output and low resistance in sepsis has also been proven: (Can Ince et al. 2018) [10], (Diamanno Ribeiro Salgado et al, 2011) [16], (A. M. Dondorp et al, 2008) [17], (Christoph Langenberg et al, 2006) [18], (Carolina Ruiz1 et al, 2010) [20], (Professor Emeritus John E et al, 2006) [22].

Animal experiments of Class I on sheep show a correlation between macro-hemodynamics and micro-hemodynamics. Therefore, the conclusion is as follows: As much of the literature has proven that the high output and low resistance in macro-hemodynamics is a universal phenomenon, the phenomenon of hyperdynamic blood flow in micro-hemodynamics is also a universal phenomenon.

(3) Proof of physiological theory: From the angle of physiological theory, the universality of hyperdynamic blood flow can be proven more sufficiently. In the physiological textbooks (Zhuda Nian et al. 2018) [11], the following contents are specified: Within a unit of time, venous returned volume is equal to cardiac output; the venous returned volume means the volume of blood flow from the vein into the right atrium in every minute; the venous returned volume and cardiac output must be equal. In the American physiological textbooks ([31] Arthur C. Guyton et al, 2016), the same viewpoints are also expressed: Venous return is the quantity of blood flowing from the veins into the right atrium each minute; the venous return and the cardiac output must equal.

Another literature report [5] specifies as follows: blood circulation in the human body is a closed loop; cardiac ejection volume is equal to venous returned volume; and thus, the cardiac output at physiological state is completely determined by venous returned volume. Therefore, hyperdynamic blood flow in micro-hemodynamics is of dynamic balance with the cardiac output. When micro-circulating massive hyperdynamic blood flow suddenly rushes into the vein, cardiac output is necessarily increased; even though the proof is not from animal experiments on sheep. Since the phenomenon of blood flow acceleration was found in the above animal experiments, it can be immediately inferred that cardiac output is necessarily increased to cause the phenomenon of high output and low resistance. To be restated, as found in the images, sublingual micro-circulating hyperdynamic blood flow mostly occurred in micro veins, because the blood in them all flows from multiple branches to a single branch; as found in animal experiments, the hyperdynamic blood flow also mostly occurred in micro vein, so that a massive blood flow to the micro veins appears. According to the physiological theory, cardiac output is necessarily increased when blood flow volume in microvein shows a large increase, because this is required for maintaining a dynamic balance in the blood system. Therefore, we reversely inferred the phenomenon of high output and low resistance, along with the physiological principle that the venous returned volume must be equal to cardiac output; it can be proven that micro-circulating massive hyperdynamic blood flow occurs before the phenomenon of high output and low resistance in macro-hemodynamics.

Although only an increase of cardiac output in sepsis was found by us in the past, it can be inferred according to the above physiologic theory that the micro-circulating hyperdynamic blood flow necessarily first occurs before and during the cardiac output increase.

As proven above according to physiological theory, hyperdynamic blood flow in micro-hemodynamics causes a high output and low resistance in macro-hemodynamics of sepsis. Therefore, when we realize that the high output and low resistance is universal, the hyperdynamic blood flow is also necessarily universal.

(4) Proof for universality from shock theory: Then, through the angle of shock classification and the heterogeneous phenomenon of micro-circulating blood flow in infective shock, the universality of hyperdynamic blood flow was proven again.

Due to the phenomenon of warm shock in distributive shock and the heterogeneous phenomenon of micro-circulating blood flow repeatedly revealed in the literature (Liuda Wei et al, 2013) [5], (De Backer et al, 2007) [4], (Can Ince at all.2018) [10], (Can Ince et al, 2015) [13] and (Bakker J et al, 2021) [27], the universality of hyperdynamic blood flow in early sepsis can be further proven.

In 1975, Weil et al. proposed a new method for shock classification according to hemodynamic characteristics: hypovolemic shock, cardiogenic shock, distributive shock and obstructive shock. These types of shock vary in treatment, and cover nearly all of clinical shock from the angle of hemodynamics. As found by Weil et al, different types of shock had different hemodynamic characteristics (Liuda Wei et al, 2013) [5].

The distributive shock is further classified into infective shock and neurogenic shock (which is caused by anesthetics overdose or nervous injury such as ganglion block and spinal shock), which varies in abnormal blood flow distribution.

See Table 2 in Page 190 of literature (Liuda Wei et al, 2013) [5].

**Table 2.** Clinical parameters and measurable related parameters of shock

| Classification | Skin | Right heart filling pressure | Cardiac output | Left ventricular pressure | Vascular resistance | Myocardial oxygen consumption |
| --- | --- | --- | --- | --- | --- | --- |
| Hypovolemia | Cold, weak | ↓ | ↓ | ↓ | ↑ | ↓ |
| Cardiogenic | Cold, weak | ↑ | ↓ | ↑ | ↑ | ↓ |
| Infectious (early stage) | Warm, red | ↔ | ↑ | ↓ | ↓ | ↑ |
| Infectious (late stage) | Cold, weak | ↓ | ↓ | ↓ | ↑ | ↓ |
| spinal | Cold, Red under the injury level | ↓ | ↓ | ↓ | ↓ | ↔ |
| Obstructive | Cold, weak | ↑ | ↓ | ↓ | ↑ | ↓ |

As shown in Table 2, the characteristics of warm skin (early stage of warm shock) was indicated only by infective shock and even not by neurogenic shock (spinal shock) which is another type of distributive shock.

From the angle of hyperdynamic blood flow, the logic of Answer 1 is as follows: the abnormal blood flow distribution (warm skin in early period and wet cold skin in late period) shown by infective type of distributive shock and the blood flow heterogeneity reflected in the literature (De Backer et al, 2007) [4], (Liuda Wei et al, 2013) [5], (Can Ince et al, 2018) [10], (Can Ince et al, 2015) [13] and (Bakker J et al, 2021) [27] are actually two fragment processes at different stages of hyperdynamic blood flow in sepsis (i.e. occurrence, development and gradual disappearance); they both reflect the same phenomenon.

From the angle of hyperdynamic blood flow, we can explain how to form an abnormal blood flow distribution or heterogeneous phenomenon in an infective distributive shock. When sepsis starts to occur, numerous toxins enter blood, and human body initiates hyperdynamic blood flow to start detoxifying. Since hyperdynamic blood flow is gradually developed at this process, some blood flow is still normal during in this period; during the occurrence of this phenomenon, this is the first type of abnormal blood flow distribution in early sepsis, i.e. warm shock of high output and low resistance of blood flow or heterogeneous phenomenon of blood flow. When the sepsis is developed into the early and middle period, all blood vessels in micro-circulating are full of hyperdynamic blood flow; strictly speaking, the abnormal blood flow distribution or heterogeneous phenomenon does not exist at this stage; because the flow rate is basically all from hyperdynamic blood flow but warm shock still exists. When the sepsis enters middle-late period, due to the long-time lack of oxygen and nutrients, some of hyperdynamic blood flow are attenuated and then gradually substituted by stagnant blood flow; some of hyperdynamic blood flow still maintain the accelerated detoxification; at this stage, typical characteristics of abnormal blood flow distribution in distributive shock or the heterogeneous phenomenon of micro-circulating blood flow occurs again; this is the second type of distributive shock or heterogeneous phenomenon in the middle and late period (i.e. cold shock of low output and high resistance). To summarize, Both of these situations are a local process throughout the entire development of hyperdynamic blood flow in sepsis.. Therefore, Answer 1 explains the characteristics of distributive shock in infective shock at sepsis or the reasons for generation of heterogeneous phenomenon.

There are two states of distributive shock in infective shock. At the first state (i.e. high output and low resistance during the early stage), the hyperdynamic blood flow is gradually expanded; in this period, normal blood flow is still maintained inside some of blood vessels without conversion into hyperdynamic blood flow; this is the first type of abnormal blood flow distribution or is called as warm shock and heterogeneous phenomenon. At the second state (in middle-late period; low output and high resistance), the hyperdynamic blood flow gradually disappears and is attenuated; in this period, some or most of hyperdynamic blood flow are converted into stagnant blood flow, but some of blood flow still keep at a state of hyperdynamic blood flow; this is the second type of abnormal blood flow distribution or called as cold shock and heterogeneous phenomenon. As a difference of these two types of abnormal blood flow distribution, the stagnant blood flow does not coexist at abnormal blood flow distribution during the early period.

After the understanding of this process and by combining the medical consensus on distributive shock and blood flow heterogeneity universally accepted by literatures, a further understanding will be made for the necessity and universality of hyperdynamic blood flow stated in answer 1. As the medical community has realized, distributive shock (including: early warm shock) is a universal phenomenon, and hyperdynamic blood flow is also a universal phenomenon. Therefore, the universality of hyperdynamic blood flow can be verified through the concept of distributive shock which has been generally accepted in medical circle.

The warm type of distributive shock is characterized by high output and low resistance; as already proven by the above physiological theory, the phenomenon of high output and low resistance is actually caused by micro-circulating hyperdynamic blood flow. Therefore, as distributive shock is a universal phenomenon of shock, the hyperdynamic blood flow is also a universal phenomenon.

In our study, the necessity and universality of hyperdynamic blood flow was first proven for the following reasons: if the hyperdynamic blood flow is proven as necessary and universal, it very probably becomes an independent influencing factor for sepsis, which is worth of deep research.

However, in historic guidelines of surviving sepsis campaign (SSC), sepsis is defined as cardiac output >3.5 L/min/m2 in 2001 guideline; but the cardiac output measurement was discontinues following formal guideline updates issued in 2002. The possible reasons are given as follows: As an invasive or semi-invasive measurement method, the cardiac output measurement is not suitable for clinical measurement of early sepsis; in addition, since such phenomenon occurred during the early period, has unobvious characteristics and is usually difficult to perceive clinically, it is often be neglected in early sepsis.

2. Question 2: Why is the blood flow acceleration (or hyperdynamic blood flow) generated? Answer 2 of the “detoxifying principle” and Proof 2 are given.

In animal experiments of Class II on rats, Question 2 is raised: Why is the blood flow acceleration (or hyperdynamic blood flow) generated?

Answer 2 of “detoxifying principle”: The blood flow acceleration (or hyperdynamic blood flow) is a detoxifying process of the human body (i.e. sympathetic-immune defense process). When the toxins in the blood exceed the allowable rang.e of the blood, hyperdynamic blood flow will be initiated to rapidly transport the toxins into relevant viscera (such as liver and kidney) for detoxifcation.

Proof 2: After the intravenous injection of LPS at reduced doses into rats, capillary blood flow started to be accelerated; meanwhile, toxic symptoms of a semi-conscious state occurred (i.e. no eating/drinking, immobile at touch and no running/jumping). After the observation for 24 hours, the activity was gradually restored to food seeking and water drinking. Three days later, normal activity was completely restored to running/jumping as usual. However, if the dose of LPS exceeded the detoxifying ability of human body, the rats would die. Since this experiment shows that a self-detoxifying function exists in the immune defense system of animal bodies, Answer 2 is established. For the detoxifying function of various organs (such as liver, kidney and lung), a clarification has been made in Chinese and foreign textbooks of “Physiology,” as well as relevant literature (Zhuda Nian et al, 2018) [11], (Professor Emeritus John E et al, 2006) [22] and (Melanie J. Scott, Timothy R et al, 2008) [25].

Literature [3] (Yaoyong Ming et al, 2018) points out that:The organism can remove and detoxify endotoxin, so that it is not easy to cause obvious damage. Liver is the main site of endotoxin inactivation and clearance, and it works through kupffer cells in monocyte/macrophage system.

And Literature [3] (Yaoyong Ming et al, 2018) At the same time pointed out that: Procalcitonin,(PCT) is a calcitonin (CT) propeptide with no hormone activity, which is a glycoprotein with a molecular weight of 13kD and composed of 116 amino acids. PCT has a half-life of 25-30 hours and good stability in vivo and in vitro … Small doses of bacterial lipopolysaccharide (LPS) injected intravenously by healthy volunteers can also induce PCT production. PCT can be detected in plasma 2 hours after LPS injection, and the concentration of PCT rises rapidly at 6-8 hours, reaching the peak at 12-48 hours, and returning to normal after 2-3 days [3]. This shows that human experiments, like our Animal (rats) experiments of Class II, also prove that the human body has the function of detoxification. It has been generally accepted and well known in the medical field. Therefore, it will not be stated any more herein.

3 Question 3: How to prove that the occurrence of accelerated (hyperdynamic blood flow) blood flow is for the purpose of “accelerating” detoxification?Answer 3 and Proof 3 are given.

For the detoxifying principle, Question 3, Answer 3 and Proof 3 are given.

Answer 3 and Proof 3 suggests that increasing the flow rate is necessary to accelerate detoxification.Logically speaking, assuming that the maximum amount of detoxification per cubic millimeter of liver cells is in the 100th percentile, the liver’s detoxification function does not need to be maximized under normal circumstances, for example, when there are very few toxins in normal blood (there are also toxins in normal blood, but they remain below normal values throughout detoxification by the liver). Once a large amount of toxins appear, high-speed blood flow rapidly transports a large amount of toxins to the liver, enabling the liver to activate its maximum detoxification ability in the 100th percentile. This high-speed blood flow plays a role in accelerating detoxification and maximizing the detoxification function of liver cells.

Therefore, it is logically proven that Answer 3 of “accelerated detoxification” is established. Such aspects are clarified in the literature (Hangyul M et al, 2008) [14]. However, this literature only shows that the blood flow acceleration can accelerate the bacterial elimination; the accelerated detoxification is not attributed to the original motivation for the human body needs for detoxifcation.

Answer 2 and 3 of “accelerated detoxification” are reasons for the generation of hyperdynamic blood flow given for the first time in the field of sepsis. Understanding the phenomenon will enable further research on the pathogenesis of sepsis. In the field of sepsis, the phenomenon of hyperdynamic blood flow has been reported in much of the foreign literature; but these literatures do not explain or understand the reasons for such phenomenon, they merely dispute the existence of hyperdynamic blood flow; this has influenced research on hyperdynamic blood flow. (De Backer et al, 2007) [4], (Vanina S et al, 2015) [8], (Vanina S et al, 2012) [9], (A. M. Dondorp et al, 2008) [17], (VS Kanoore Edul et al, 2015) [19], (Arnaldo Dubin et al, 2020) [23], (Bakker J et al, 2021) [27]. For example: the “Second Consensus on Sublingual Micro-circulating Assessment of Critical Patients” (2018, European Society of Intensive Care Medicine) specifies as follows: Although its origin and clinical significance still remains to be determined, the existence of hyperdynamic blood flow can be explained as micro-circulating variation (Can Ince et al, 2018) [10]. Its origin is determined through the principle of “accelerated detoxification” in Answer 2 and 3, which is our first step for solving the dispute on hyperdynamic blood flow.

Question 4: What adverse impact on the human body will be produced if the micro-circulating blood flow rate inside the true capillaries exceeds the normal limit? Answer 4 of “the hyperdynamic blood flow causes Xinghuai Feng-Bernoulli warm shock” and Proof 4 are given.

At present, classical theory (i.e. shunting theory) only explains the warm shock of high output and low resistance as follows: At infective or toxic shock, arteriovenous shunt and thoroughfare channels are opened in large numbers; the patients are in a state of shock, but with warmer skin (i.e. warm shock); since massive microarterial blood enters through anastomotic branch into microvein and does not make material interchange with histiocytes, tissue anoxia is aggravated to exacerbate the illness state (Zhuda Nian et al, 2018) [11]. However, this theory only gives indirect cause for warm shock, and does not directly explain whether the existence of hyperdynamic blood flow inside the true capillaries is related to warm shock.

The following diagram visually expresses the description of physiological textbooks ((Zhuda Nian et al, 2018) [11], Figure 4 on Page 124).

**Figure 4.**
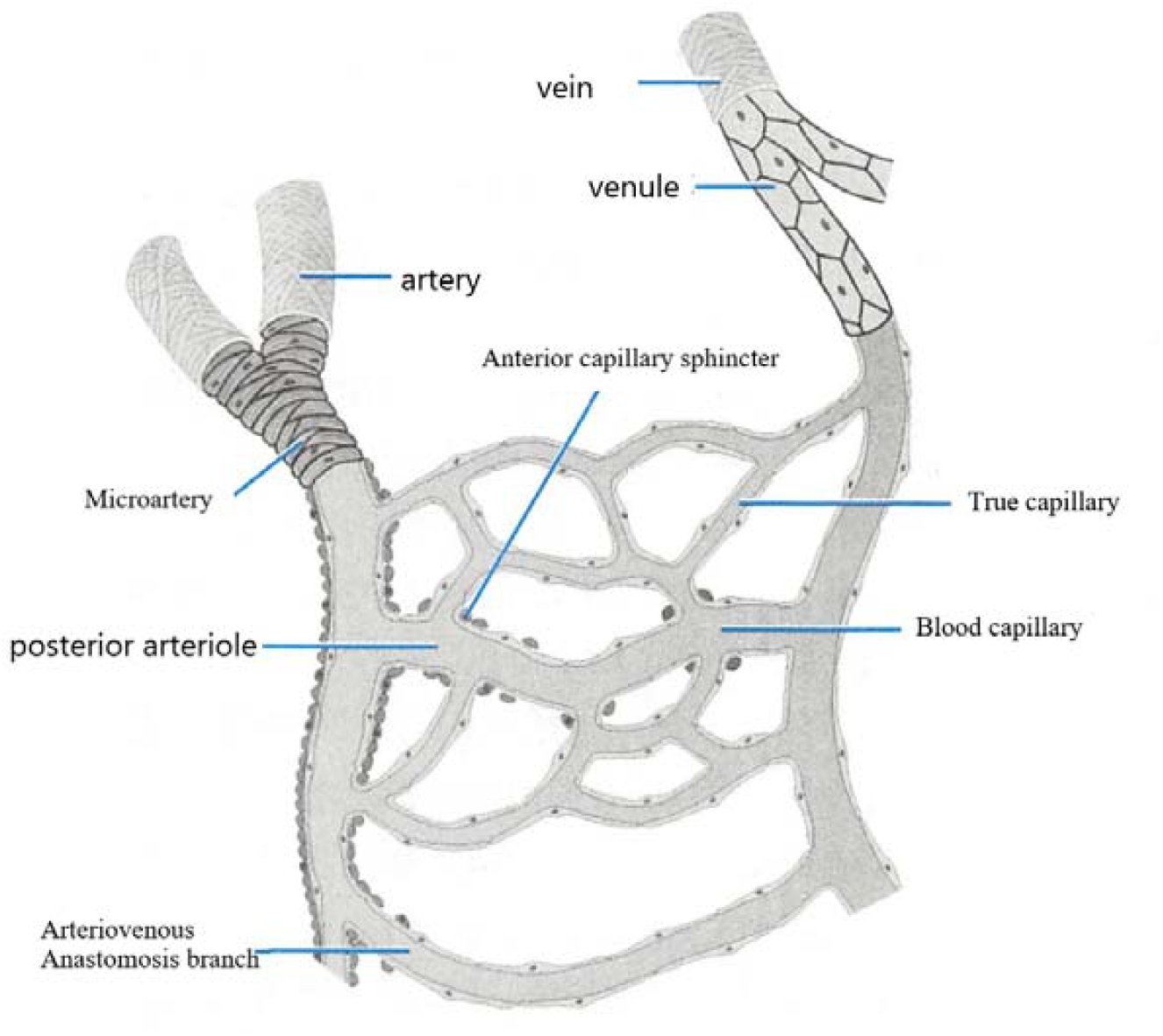
Composition pattern diagram of Microcirculation

We know that: Since the real site for oxygen exchange is the true capillary network, direct cause for oxygen exchange disorder is attributed to abnormalities inside the true capillary network. Then, there are four possible conditions for that massive microarterial blood enters through anastomotic branch into microvein and does not make material interchange with histiocytes. Condition 1 (which is clarified in above textbooks): Since microarterial blood inside true capillary network enters through anastomotic branch directly into microvein (shunting theory), it is potentially inferred that this will cause a shunting and evacuation of blood inside the true capillary network; therefore, oxygen exchange disorder does not occur in blood inside the true capillary network; under this condition, there is no blood flow inside the true capillary network, meaning that internal respiration stops and the human body immediately dies by suffocation, and there will be no continuous phenomenon of the warm shock found in clinical practice; therefore, this condition does not conform to clinical facts. Condition 2: Inside the true capillary network, some blood still flows slowly, which does not consist with the results of our animal experiments, because the blood flow is accelerated; this also does not conform to the phenomenon of hyperdynamic blood flow in warm shock found in clinical practice; logically speaking, if some blood still flows slowly, clinical manifestations should be cold shock; however, the actual clinical manifestation is warm shock. Condition 3: Inside the true capillary network, all blood is stagnant and there is no perfusion or flow, which does not consist with the phenomenon of hyperdynamic blood flow found in the above animal experiments and the results of its clinical observation (De Backer et al, 2007) [4]. Therefore, only Condition 4 is applicable: Inside the true capillary network, the blood flows at a very high rate (called hyperdynamic blood flow), which is consistent with the results of experimental observation and the manifestations of clinical observation.

Therefore, oxygen exchange disorder may only occur in hyperdynamic blood flow inside the true capillary network and microveins; its reason can be attributed to the Bernoulli principle; it is called by us the Xinghuai Feng-Bernoulli warm shock mechanism, to distinguish it from the shunting theory in current textbooks. Next, we will make a deep theoretic exploration of the Xinghuai Feng-Bernoulli warm shock mechanism. This deep analysis will not only reveal the direct cause for warm shock, but also very probably reveal the pathogenesis for sepsis.

How the Bernoulli principle becomes the direct cause for warm shock is concretely proven in the following. In order to make such proof, we have to quote a large section of knowledge from the physiological textbooks [11], the understanding of the reader is appreciated.

(1) According to physiological theory, oxygen exchange and carbon dioxide exchange in the human body is determined by a difference in pressure. The physiological textbook (Zhuda Nian et al, 2018) [11] specifies the following: Tissue ventilation is the exchange in gas between blood in systemic circulation capillary and histiocytes. Between tissue ventilation and pulmonary ventilation, the similarity lies in mechanism and influencing factors; the difference is that the gas exchange occurs between the liquid medium (i.e. blood, tissue fluid and intracellular fluid) and the partial pressure difference of O2 partial pressure (PO2) and CO2 partial pressure (PCO2) between the two sides of the diffusion membrane. This varies with the intensity of intracellular oxidative metabolism and the volume of tissue blood flow. If the blood flow volume is unchanged and the metabolism is enhanced, a PO2 decrease and PCO2 increase will occur in the tissue fluid; if metabolic rate is unchanged and the blood flow volume is increased, a PO2 increase and PCO2 decrease will occur in tissue fluid. Due to the aerobic metabolism of cells, the utilization of O2 and the generation of CO2, PO2 will be as low as <30 mmHg, and PCO2 will be as high as >50 mmHg. When arterial blood flows through tissue capillary, O2 diffuses according to partial pressure difference from blood to tissue fluid and cells; CO2 diffuses from tissue fluid and cells to blood; due to the loss of O2 and the obtaining of CO2, arterial blood becomes venous blood.

Gas molecules move continuously in an astatic way. When gas pressure differences exist between different regions, gas molecules will make a net transfer from sites with high gas pressure to those with low gas pressure, this is known as gas diffusion. According to its own partial pressure difference, each of these mixed gases diffuse from sites of high partial pressure to those with low partial pressure until a dynamic balance is reached. Both pulmonary ventilation and tissue ventilation are accomplished using diffusion. The volume of gas diffusion within unit time is defined as the gas diffusion rate (D). Fick Dispersion law specifies as follows: When gas passes through thin-layer tissue, gas diffusion rate is directly proportional to gas partial pressure difference between two sides of tissue (AP), temperature (T), diffusion area (A) and gas molecule solubility (S); but it is inversely proportional to diffusion distance (d) and square root of gas molecular weight (MW). The correlation of gas diffusion rate with each influencing factor is shown in the following formula below:

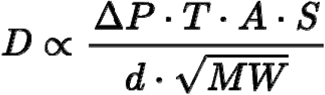

Gas partial pressure difference: Gas partial pressure means the pressure produced by each gas component of mixed gas. Under constant temperature, partial pressure of a certain gas is obtained by multiplying the total pressure of mixed gas and the volumetric ratio of this gas among mixed gas. For example: Air is a mixed gas with a total pressure of 760 mmHg, the O2 volumetric ratio is about 21% and PO2 is 760×21% = 159 mmHg; CO2 volumetric ratio is about 0.04%, and PCO2 is 760×0.04% = 0.3 mmHg. Gas partial pressure difference (□P) means the difference value in partial pressure of a certain gas between two regions; it is the dynamic force for gas diffusion and the key factor for determination of gas diffusion direction.

(2) Characteristics of Hb-O2 combination: This combining reaction is rapid (<0.01 seconds), reversible, and dissociates very rapidly. Both combination and dissociation do not require an enzymatic catalysis, but can be influenced by PO2. When blood flows through lungs with high PO2, Hb is combined with O2 to form oxyhemoglobin (HbO2); when blood flows through tissue with low PO2, HbO2 is rapidly dissociated to release O2 and become Hb. This process can be expressed through the following formula:

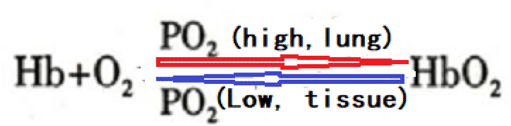

(Zhuda Nian et al, 2018) [11]. Therefore, partial oxygen pressure difference is the direct dynamic force for the oxygen exchange.

(3) Bernoulli equation:

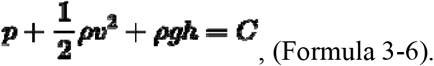

According to the Bernoulli equation, the following phenomenon is shown. When an ideal fluid flows steadily inside the flow tube, the kinetic energy in unit volume, the gravitational potential energy in unit volume and the sum of pressure intensity at the site is constant. In the Bernoulli equation, three terms possess the dimension of pressure intensity: the term 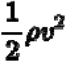 is related to the flow rate, and is often called dynamic pressure; the term p and pgh are unrelated to the flow rate; p is often called static pressure. If a fluid flows inside a horizontal tube (h1=h2), the potential energy of the fluid system is unchanged during the flowing course. Formula 3-6 can be written to Formula. 3-7: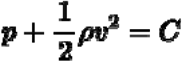.

As shown by the above formula, fluids flowing inside horizontal tube have a pressure intensity that is larger at sites with a low flow rate; and a pressure intensity that is smaller at sites with a high flow rate [34].

(4) Gas partial pressure means the pressure produced by each gas component of mixed gas. When under constant temperature, partial pressure of a certain gas is obtained by multiplying the total pressure of mixed gas and the volumetric ratio of this gas in a mixed gas [11].

(5) According to Bernoulli equation (Formula 3-6 and 3-7), the following phenomenon is shown. When blood flow rate is increased to a certain extent, the pressure intensity of arterial blood drops to reduce the oxygen partial pressure, making it difficult for oxygen molecules in arterial blood to enter tissue fluid for the oxygen supply of cells by relying on oxygen partial pressure difference, which causes oxygen exchange disorder (i.e. warm shock). This is just the basis for warm shock being caused by hyperdynamic blood flow inside true capillaries and microveins, called Xinghuai Feng-Bernoulli warm shock.

The direct cause for warm shock is explained above, through the Xinghuai Feng-Bernoulli warm shock mechanism. The current textbooks are just unilateral for the shunting theory of warm shock. This will lay a solid foundation for revealing the pathogenesis of sepsis.

In addition, the logic for Answer 4 is as follows. If the source of infective toxins exists continuously, a hyperdynamic blood flow will be continuously generated according to the principle of “accelerated detoxification” mentioned in Answer 2 and 3; at the continuous micro-circulating hyperdynamic blood flow, chronic anoxia will be caused in the human body, the clinical manifestations of which are warm shock symptoms of sepsis with high output and low resistance (Yaoyong Ming et al, 2018) [3], (Liuda Wei et al, 2013) [5], (Zhuda Nian et al, 2018) [11], (Diamantino Ribeiro Salgado et al, 2011) [16], (A. M. Dondorp et al, 2008) [17], (Carolina Ruiz1 et al, 2010) [20], (Professor Emeritus John E at all.2006) [22].

If intervention and rescue are not implemented, the continuous warm shock will cause anoxia and nutrient deficiency of cells/organs, which will start to fail. Then, the hyperdynamic blood flow is also gradually attenuated, and the stagnant blood flow is gradually increased correspondingly,I. e. the beginning of “cold shock”; this is micro-circulating blood flow heterogeneity as revealed in the literature (Paul WG Elbers et al, 2006) [6], (Vanina S et al, 2015) [8], (Vanina S et al, 2012) [9] and (Can Ince et al, 2018) [10]. Finally, this state transitions to cold shock, subsequent disseminated intravascular coagulation (DIC) and multiple organ dysfunction syndrome (MODS).

If the mechanism of accelerated detoxification in Answer 2 and 3 explains the principle of immune defense function in human body and provides a train of causal thoughts for generation of hyperdynamic blood flow, the Xinghuai Feng-Bernoulli warm shock mechanism explains the direct pathogenesis of hyperdynamic blood flow for how to cause oxygen exchange disorder and thus induce warm shock in early sepsis,And then it also explained the cause of cold shock.

Answer 2 and 3 states: The generation of hyperdynamic blood flow is a spontaneous compensatory reaction of human body (i.e. detoxifying effect). This has a certain advantages for human body. However, since any matter is double-sided, the hyperdynamic blood flow also has side effects unfavorable for human body. We utilize the Xinghuai Feng-Bernoulli warm shock mechanism to explain the disadvantages of hyperdynamic blood flow that oxygen exchange disorder is caused to induce warm shock. This mechanism may be greatly significant not only from various angles (such as exploration of pathogenesis and finding of medical science) but also for guidance on clinical work and research work.

In fact, the oxygen exchange between blood and cells is influenced by many factors, such as erythrocyte deformability, capillary density, hematocrit and blood flow rate. However, we should find out main influencing factors oxygen exchange under particular conditions. When in a normal state, the above factors can influence the oxygen exchange; under the state of hyperdynamic blood flow, they can not become main influencing factors because the difference is too large between internal and external pressure of the capillaries, but the blood flow acceleration beyond normal limits will become main influencing factors. Of course, hyperdynamic blood flow can be considered as a main influencing factor for oxygen exchange only under the premise that the effect of the Bernoulli principle should be understood and can be utilized to explain the reasons for oxygen exchange disorder (i.e. warm shock). If the Bernoulli principle is not understood, there is no way to grasp this main factor. For example: In the literature (A. M. Dondorp et al, 2008) [17], it is considered that the hyperdynamic blood flow can not cause an oxygen exchange disorder.

Question 5: Where does the direct dynamic force for hyperdynamic blood flow come from? Answer 5 of the “pulling principle” and Proof 5 are given.

Regarding the source of direct dynamic force for hyperdynamic blood flow, the physiological textbook (Zhuda Nian et al, 2018) [11] proposes the following: in infective or toxic shock, arteriovenous shunt and thoroughfare channels are opened in large number. This can illuminate our research.

Answer 5: In early sepsis, micro-circulating thoroughfare channels and arteriovenous anastomotic branch are opened in large number to pull the blood flow acceleration to the microveins (which is called the pulling principle). Proof 5: In fact, this is also an embodiment of the Bernoulli principle. After the micro-circulating arteriovenous anastomotic branch is opened, a negative pressure is formed for the blood flow inside the true capillary exchange network and microveins to produce a sucking effect. See the following Figure 5:

**Figure 5:**
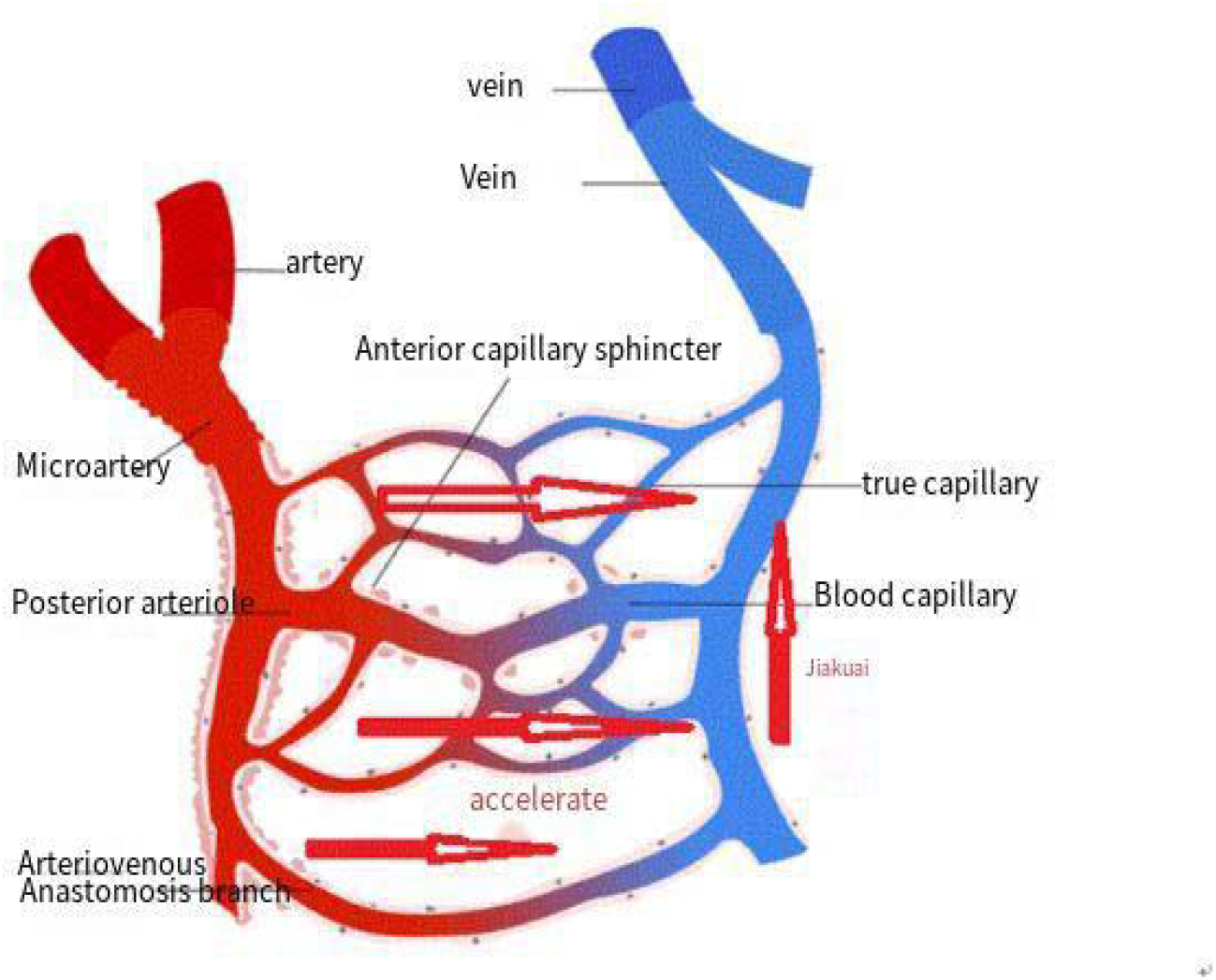
After a short circuit in the blood flow of the anastomotic branch, the accelerated flow pulls the blood flow of the true capillaries, generating hyperdynamic blood flow.

Therefore, the shunting theory in physiological textbooks states the following: Due to shunting of the anastomotic branch, some blood can not participate in oxygen exchange, meaning it cannot become a main or direct cause. Since blood flow exists in true capillaries, the main and direct cause for oxygen exchange disorder is abnormalities in the true capillaries (i.e. occurrence of hyperdynamic blood flow).

Conclusion: According to the above studies, we propose the following pathogenesis of sepsis.

Through overall consideration of the above five questions and the corresponding answers and proofs, we clarify the pathogenesis of sepsis as integrally as possible from the angle of micro-circulating hyperdynamic blood flow. At the beginning, the toxins invade from wounds into the blood system; when numerous toxins are found in blood, the human body will begin hyperdynamic blood flow to rapidly transport the toxins to relevant organs (such as liver and kidney) for detoxifcation. As side effects of this process, blood flow inside the true capillaries is so rapid that it exceeds the normal limit of micro-circulating blood flow rate in human body (i.e. V>1000μm/s, Attachment II). According to the Bernoulli principle, in capillaries with blood flow acceleration, the pressure is necessarily decreased to reduce the oxygen partial pressure inside the blood vessels and decrease the difference between internal and external oxygen partial pressure in the capillary walls, so that most oxygen molecules have difficulty in making a normal diffusion outside the blood vessels for the oxygen supply of cells, which causes oxygen exchange disorder. Due to the effect of detoxifying principle, the toxins continuously invade from the wound into the blood system usually, within a few days. Due to the pulling principle, hyperdynamic blood flow continuously occurs in the human body under the action of the immune defense system, which causes Xinghuai Feng-Bernoulli warm shock. The occurrence and development of shock is a continuous process: presenting as warm skin and basically normal blood pressure in Xinghuai Feng-Bernoulli warm shock; and wet, cold skin, piebald formation and blood pressure decrease in cold shock. If timely treatment is not given at the stage of warm shock, cells and organs cannot obtain oxygen and nutrients as usual over a period of time; in continuous warm shock, hyperdynamic blood flow is gradually attenuated, and stagnant blood flow is gradually increased; the human body will gradually enter the stage of cold shock, with wet, cold skin and a blood pressure decrease, and typical symptoms of septic shock will occur (i.e. typical manifestations of infective type of distributive shock) If effective treatment is not yet given, DIC and MODS will ultimately occur. From the angle of micro-circulating hyperdynamic blood flow, the whole development process of pathogenesis of sepsis is explained. As the cause for warm shock, cold shock and MODS, the hyperdynamic blood flow in sepsis very probably becomes main pathogenesis of sepsis.

Our theory on the pathogenesis of sepsis also answers the following query of literature (Can Ince et al, 2018) [10]: Although its origin and clinical significance still remain to be determined, the existence of hyperdynamic blood flow can be explained as micro-circulating variation.

Sepsis is defined as a life-threatening organ dysfunction caused by host-response disorder under infective conditions (Mervyn Singer et al, 2016) [2]. The theory for sepsis pathogenesis through hyperdynamic blood flow meets three elements of this definition: infection, host response disorder and life-threatening organ dysfunction. Moreover, this theory clarifies the concrete organs and phenomenon of host response disorder: firstly, the host is just the blood flow inside micro-circulating true capillaries; secondly, in host response disorder (i.e. hyperdynamic blood flow occurs inside true capillary for a long time), internal environment steady state of blood flow is destroyed to cause oxygen exchange disorder (i.e. Bernoulli warm shock); finally, the continuation of anoxia in warm shock causes cold shock to induce MODS.

From the angle of hyperdynamic blood flow, we propose the theory of sepsis pathogenesis. There are two foremost innovative basic principles: one is the detoxifying principle; and the other is Xinghuai Feng-Bernoulli warm shock mechanism. These two innovative principles are the cornerstone for pathogenesis of sepsis.

In order to correctly find hyperdynamic blood flow, a reference should be made to Attachment III “Inclusion criteria for hyperdynamic blood flow sample of clinical sepsis, suggestions/precautions for sampling method and limitations of common space-time method in measurement of hyperdynamic blood flow”

Potential clinical significance of theory for sepsis pathogenesis

As is already known, the sepsis positive rate is very low in blood culture tests in clinical practice (Lin-hong Yuan, J.2018 et al) [28]. Blood culture as the golden standard requires a long time to find pathogens; up to 70% of sepsis patients have received anti-infective treatment; so as to influence the finding time and positive rate of pathogen. Therefore, it possesses a very important clinical value to seek for more rapid specific indices for clinical diagnosis of sepsis (Lu-qiu Wei, et al) [35].

Firstly, we should realize that the human body can be compared to a highly-sensitive and accurate biological laboratory. It possesses a very high accuracy for finding of various pathogens for the following reasons: through the evolution of more than 0.1 billion years, the human body possesses a very strong ability for pathogen identification; otherwise, human beings will be die off at a young age by nature according to the law of survival of the fittest. Now, to determine if the test results of this medical laboratory and the immune defense process for initiation of hyperdynamic blood flow inside human body; particularly, the correct understanding of this process will bring about very good potential practical indices for clinical practice. We realize that: the human body is a highly-sensitive and accurate laboratory; by utilizing this laboratory, sepsis can be accurately and specifically diagnosed early.

Clinical value 1: Early and ultra early detection of sepsis, i.e. when the hyperdynamic blood flow of sepsis has just occurred.

Clinical value 2: As specified in “International Consensus on Sepsis (3.0)” (Mervyn Singer et al, 2016) [2], work groups have attempted to differentiate the sepsis from simple inflammation. Such differentiation is very simple, Because simple inflammation does not cause the appearance of hyperdynamic blood flow. For example: Among two inflammation patients, sublingual hyperdynamic blood flow occurs in one patient, but micro-circulating is normal in the other patient; the former develops into sepsis, but the latter is simple inflammation.

Clinical value 3: Hyperdynamic blood flow is utilized for accurate judgment of sepsis in the early, middle and late period, so as to make a scientific judgment of prognosis and take different measures for rescue/treatment. Manifestations in different periods of sepsis have been stated above: In the early period, hyperdynamic blood flow occurs to indicates that the human body will begin the detoxifying process; in the middle-late period, hyperdynamic blood flow starts to be attenuated and substituted by stagnant blood flow; in the late period, there is necessarily no blood perfusion/flow. In the literature (Vanina S et al, 2015) [7], hyperdynamic blood flow was not found, possibly because the sepsis was not staged. The literature (Zhangxiao Lei et al, 2021) [29] and (Geri et al, 2019) [30] state as follows: as compared with that with hyperdynamic blood flow, 30-day cumulative survival rate was lower in patients with stagnant and dilutive blood flow, and the difference was statistically significant; indicating that the hyperdynamic blood flow only occurs in early sepsis. In the middle-late period and late period, the stagnancy and no perfusion/flow of blood is a necessary physiological phenomenon. As compared with the stagnancy and no perfusion/flow of blood, hyperdynamic blood flow only occurs in early period. When hyperdynamic blood flow starts to be attenuated, some blood flow becomes stagnant; the other blood still maintains the state of hyperdynamic blood flow; this is middle period. Therefore, this is logically a necessary conclusion.

Clinical value 4: The international requirement is realized that the duration of bundle treatment should be shortened from 3 hours to 1 hour. When numerous toxins occur in blood, the human body starts hyperdynamic blood flow for detoxification. Once this process is found, various preparatory work can be made in advance and even at shorter time than 1 hour required for bundle treatment (In fact, we have already predicted sepsis 12 to 24 hours ahead of schedule, providing more ample preparation time for cluster therapy (clinical cases are expected to be published next year).

Secondly, as found in our study, there are two forms of hyperdynamic blood flow. The first form,Since this behavior is akin to a waterfall, therefore,the hyperdynamic blood flow in microvessels with a diameter of 50-100 μm is called “waterfall blood flow”.The second form,Since this behavior is akin to a swarm of flying mosquitos, the hyperdynamic blood flow in thinner blood vessels with diameter of <20 μm is called “flying mosquitos blood flow”. Such denomination can facilitate a rapid identification of hyperdynamic blood flow in clinical practice.

In “International Consensus on Sepsis (3.0)” (Mervyn Singer et al, 2016) [2], the sepsis defined as the life-threatening organ dysfunction is caused by host-response disorder at infection; this new definition emphasizes that the host response disorder plays a crucial role in infection. Although such judgment is correct, concrete manifestations of host-response disorder are yet unknown. There is an urgent clinical need to know the manifestations of host-response disorder, so as to apply them for clinical guidance in life-saving care. As shown by our study, the host should be the blood flow inside true capillary network; during host-response disorder, micro-circulating hyperdynamic blood flow exceeds the upper limit of normal value in human body to cause Bernoulli warm shock; therefore, the micro-circulating hyperdynamic blood flow was exactly the concrete forms and characteristics of host response disorder. This solves the query mentioned in “International Consensus on Sepsis (3.0)”: “The task force recognized that no current clinical measures reflect the concept of a dysregulated host response”.

Finally, an analysis was made for the relation between the current knowledge on sepsis pathogenesis in Chinese and foreign medical circle and the viewpoints in our study. It took a long time for humans to discover sepsis. The international definition of sepsis has changed from Version 1.0 in 1991 to Version 3.0 in 2016. The field for sepsis pathogenesis has also expanded to frontier scientific fields (such as cytobiology, genetics, immunology, molecular biology and genomics) (Yaoyong Ming et al, 2018) [3]. In early sepsis, pro-inflammatory and anti-inflammatory reactions occur; the main change of non-immune channels are complicated (such as cardiovascular system, nerve, autoregulation, endocrine, biological energy, metabolism and blood coagulation); there are multichannel molecular characteristics (such as transcription, metabonomics and proteomics) (Mervyn Singer et al, 2016) [2]. The pathogenesis and manifestations of sepsis have been discovered from various aspects: bacterial pathogenic factors; effects of inflammatory mediator; relations between endothelial cell injury (permeability of endothelial cells) and micro-circulating disorder; systemic inflammatory reactions; blood coagulation dysfunction; genetic polymorphism; high metabolism; mitochondrion oxygen utilization disorder; disseminated intravascular coagulation; apoptosis; immunosuppression and apoptosis; enteric bacteria and bacterial endotoxins (LPS) translocation; neuro-endocrino-immune network) (Yaoyong Ming et al, 2018) [3], (Fu Yuan et al, 2014) [12], (Linhong Yuan et al, 2018) [28]. However, the primary and main cause should attract attention. As shown by our study and clinical literatures (Yaoyong Ming et al, 2018) [3], ((Liuda Wei et al, 2013)) [5] and (Zhangxiao Lei et al, 2021) [29], the chronic anoxic state caused by hyperdynamic blood flow accompanied the body from early period of sepsis (3 minutes after LPS injection) until the death. During the developing course from hyperdynamic blood flow to stagnant blood flow and from warm shock to cold shock, the anoxic state is probably the primary and main cause among numerous mechanisms for sepsis pathogenesis. Due to long-time anoxia, normal immune defense function in human body declines to cause the occurrence of a series of disorders.

For example: If a mammal is suffocated by airway obstruction, it will necessarily die within a short time. During this course, every internal steady-state system of body will necessarily be destroyed. If the above pathogenesis is detected in time, a series of abnormalities will necessarily be found in terms of physiology, pathology and biochemistry (Mervyn Singer et al, 2016) [2], such as the variation at the level of cell and molecular structure. However, We should not forget that the primary and main cause is the anoxia caused by airway obstruction.

In scientific research, changing the angle of study is one way to further understanding of a problem. Through the research sepsis from the angle of hyperdynamic blood flow, the mysterious pathogenesis of sepsis may ultimately be revealed.

## Conclusion

This study systematically elucidated the occurrence and progression of sepsis in the absence of complications through the observation of blood flow velocity and analysis of underlying principles. It starts with the body detecting the invasion of a large amount of toxins into the bloodstream - the “body laboratory recognition principle”. To detoxify as quickly as possible, the body initiates high-speed microcirculatory blood flow - the “detoxification principle”. Unfortunately, the “Xinghuai Feng-Bernoulli warm shock principle” causes a decrease in oxygen partial pressure leading to warm shock. If left untreated, warm shock continues to progress and forms cold shock, where microcirculatory blood stasis occurs. Cold shock leads to more severe hypoxia, which ultimately results in multiple organ dysfunction and failure due to hypoxia in various organs. This reaches the final stage, which is sepsis as defined in Sepsis 3.0: “Sepsis is defined as life-threatening organ dysfunction caused by a dysregulated host response to infection”. The primary pathogenic mechanism of early sepsis is hypoxia caused by “Xinghuai Feng-Bernoulli warm shock”.

This article was completed in June 2023, and it has been over a year since then. During this period, we collaborated with three Class A tertiary hospital to conduct clinical observations and verifications of the principles outlined in this article. The verification was divided into severe, mild, and normal groups. Severe patients were validated with infection + SOFA >= 2, while mild patients were observed with infection as the target, referring to SOFA data. In total, there were approximately more than 500 cases, and the compliance rate was very high. The specifics will be published in a joint article by the three hospitals. Here, we only illustrate that the principles of this article have undergone multicenter, randomized, and double-blind clinical verification and have achieved satisfactory results.

## Acknowledgements

We thank Dr. Zhang Peng from the Affiliated Hospital of Qingdao University for his help with animal experiments.

## Author contributions

All authors who meet the criteria for authorship are included in the list of authors. Each author has confirmed their participation in the conception, design, analysis, writing, or revision of the manuscript. They have also contributed to the analysis and interpretation of results within their respective domains of interest. Xinghuai Xinghuai Feng conceived and directed the study, as well as conducted the majority of the experiments. Y. Zeng performed animal experiments and provided theoretical evidence regarding the relationship between macroscopic hemodynamics and microcirculation hyperdynamic blood flow. Y.B. Sun conducted animal experiments and collected and analyzed data. Bu-Wei Yu reviewed and revised the article. Xinghuai Feng and Wei Liu prepared the initial draft of the manuscript and coordinated its finalization. All authors have approved the final version of the manuscript.

## Availability of data and materials

The data sets generated and/or analyzed during the current study are not publicly available but are available from the corresponding author on reasonable request.

## Declarations

### Ethics approval and consent to participate

This study was approved by the Animal Care and Use Committee of the Sichuan University and Qingdao University. All animal experiments conformed to relevant stipulations of Chinese governments on welfare and ethics of experimental animals(GB/T 35892-2018).

### Consent for publication

Not applicable.

### Author details

1 Xuzhou Lihua Electronic Technology Development Co., Ltd.2 Department of Hypertension and Endocrinology, Daping Hospital, Chongqing, China.3 Affiliated Hospital of Qingdao University.4 West China School of Basic Medical Sciences & Forensic Medicine, Sichuan University.5 Ruijin Hospital Affiliated to Shanghai Jiaotong University School of Medicine

## Attachment I

The following two pictures are sublingual microcirculation videos for one case (cc6118-S3) of severe sepsis which are published in the attachment [Additional file 3] of literature (Daniel De Backer, et al 2007) [4]. Within the visual field, the highest flow rate was about 3307 μm/s. is shown in Figure 6:

**Figure 6:**
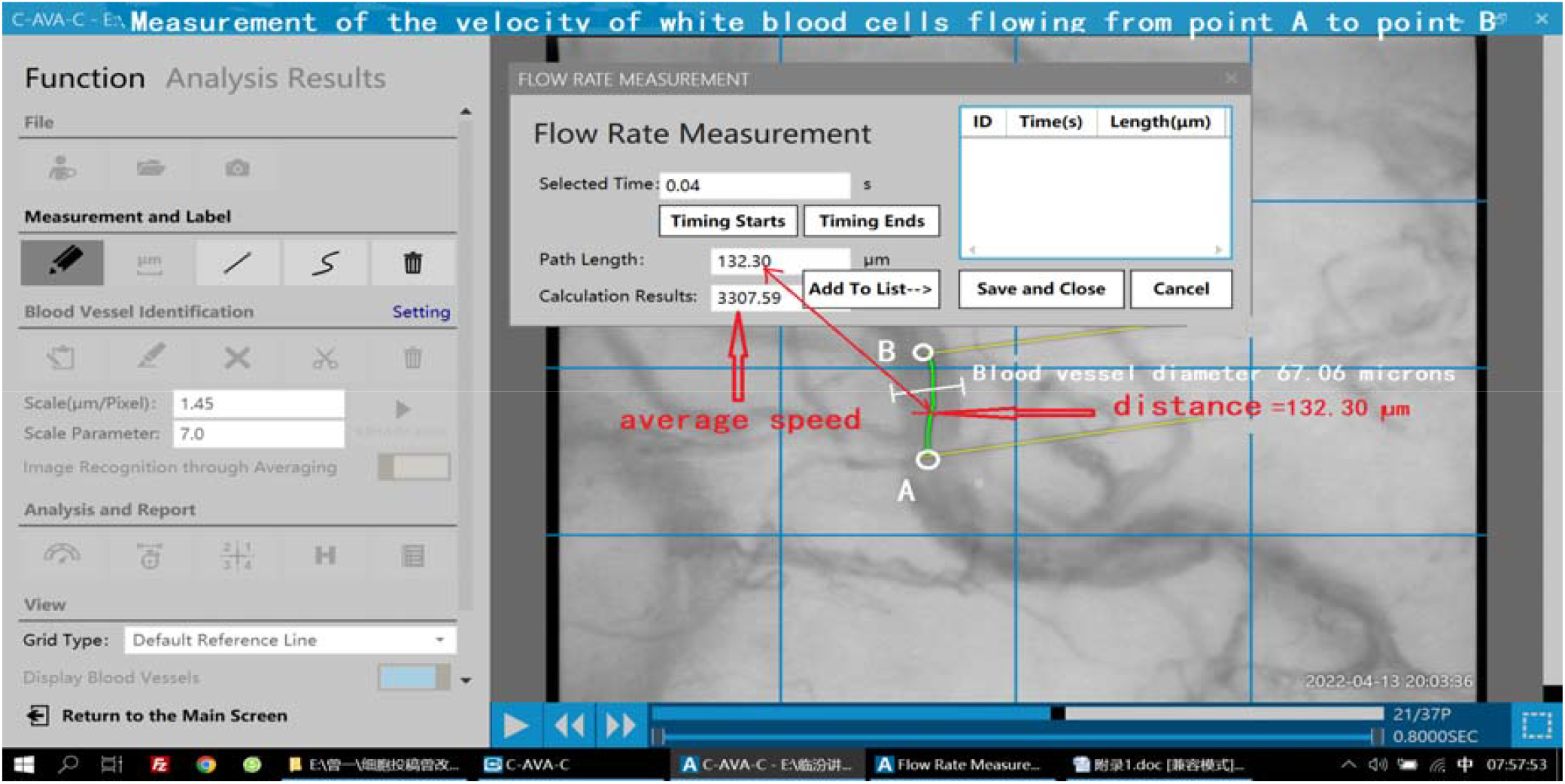
In capillary with diameter of 67.06 μm, the velocity of hyperdynamic blood flow was 3307.59 μm/s; the flowing distance of leukocyte from Frame 21 to Frame 22 was 132.30 μm.

Measurement method for blood flow rate: Through the macroscopic comparison, one blood vessel with highest blood flow rate is first selected as test sample (vascular diameter is not considered; the principle is shown hereafter). Then, the motion path of one erythrocyte/leukocyte in this blood vessel is traced. The flowing distance of erythrocyte/leukocyte from Point A to Point B is calculated through computer; the flowing time at this path is calculated through computer. Finally, the distance is divided by the time to obtain mean blood flow rate.

### Note

1. Principle for that vascular diameter is not considered: According to the literature (Liuda Wei et al, 2013)[5],blood flow rate is 500-1000 μm/s in normal people; threshold value of hyperdynamic blood flow is >1000 μm/s. At quiescent state, normal blood flow rate in microcirculation is ≤1000 μm/s. Therefore, the blood vessel is selected not by considering the diameter but according to the highest blood flow rate. Only the blood flow inside this blood vessel may cause Xinghuai Feng-Bernoulli warm shock. See the text and Attachment II.
2. In the microvessel with diameter of <20 μm, there is an interference of pulse blood; and the pulse blood is easily mistaken as hyperdynamic blood flow. Therefore, it is not suggested to measure the blood flow rate in microvessel with diameter of <20 μm. On the contrary, in the blood vessel with diameter of >20 μm, there is no interference of pulse blood; and the blood flow rate is stable. If blood flow is accelerated in this blood vessel, an abnormal variation can be considered.

## Attachment II

We studied the source for conclusion of literature(Liuda Wei et al, 2013) [5] that normal blood flow rate in capillary is 500∼1000 μm/s. Its theoretical basis is is shown in the literature (Feng Yuanzhen, *et al*.,1986)[21], (Professor Emeritus John E et al, 2006) [22]. From physical angle, the motion law of blood flow should necessarily conform to the conservation law of mass, momentum and energy. For any given spatial region, the conservation law of mass implies as follows: any inflowing matter is necessary to flow out (Feng Yuanzhen (USA),*et al*.,1986) [21]; according to the conservation law of momentum, blood flow rate is inversely proportional to vascular cross-sectional area. The blood flows from aorta through medium artery and microartery to capillary, and then from microvein to return through caval vein to heart; despite of large aperture, there is only one piece of aorta; despite of small aperture, there are innumerable pieces of capillary. Therefore, total cross-sectional area of capillary is about 220∼440 folds larger than the cross-sectional area of aorta. Mean linear velocity of blood flow in aorta is about 220 mm/s; as calculated through above formula, mean linear velocity of blood flow in capillary should be 220/440∼220/220 of this value (i.e. 0.5∼1.0 mm/s), which basically consists with the measured value. Therefore, blood flow rate >1000 μm/s is considered as abnormal state which exceeds the physiological upper limit of normal blood flow rate in microcirculation, i.e. initial threshold value of the velocity of hyperdynamic blood flow (V). In Page 577 of literature [5], the following contents are stated: At calm state, mean blood flow rate inside aorta of human being is 180∼200 mm/s, and mean blood flow rate inside capillary is 0.3∼1.0 mm/s; and there is a difference of 200∼600 folds. The literature (Professor Emeritus John E et al, 2006) [22] states as follows: Blood flow rate is inversely proportional to vascular cross-sectional area. Therefore, at quiescent state, mean blood flow rate inside aorta is about 33 cm/s; mean blood flow rate inside capillary is 1/1000 of this value, i.e. about 0.3 mm/s. In all of above literatures, the range of blood flow rate inside capillary is inferred according to the conservation law of mass (any inflowing matter is necessary to flow out) and the conservation law of momentum (the velocity is inversely proportional to area), which consists with the measured value.

Above contents are the basis for that the capillary blood flow rate of 1000 μm/s is determined as initial threshold value of the velocity of hyperdynamic blood flow. In order to avoid a confusion, the intension for this initial threshold value is strictly specified as follows:

1. The microcirculating hyperdynamic blood flow in sepsis is the manifestations of microcirculation at quiescent state of human body under the phenomenon of high output and low resistance in macro-hemodynamics.
2. Concretely, the microcirculating hyperdynamic blood flow means the blood flow with velocity of >1000 μm/s inside true capillary exchange network and microvein at microcirculating capillary circuitous channel; and this velocity is just the initial threshold value of microcirculating hyperdynamic blood flow in sepsis. For the warm shock high output and low resistance, the current classical theory (i.e. shunting theory) explains as follows: At infective or toxic shock, arteriovenous shunt and thoroughfare channel is opened in large number; the patients are at shock state but of warmer skin (i.e. warm shock); since massive microarterial blood enters through anastomotic branch into microvein and does not make material interchange with histiocytes, tissue anoxia is aggravated to exacerbate the illness state (Zhuda Nian et al, 2018)[11]. However, as known through the analysis, since oxygen exchange is really made inside true capillary network, oxygen exchange disorder is essentially an abnormality in true capillary network. Therefore, the classical shunting theory is just unilateral for explaining the warm shock of high output and low resistance; because it does not realize that the real place for oxygen exchange is inside true capillary network. Therefore, real cause for warm shock is the abnormality of blood flow inside true capillary network, but is not from the shunting theory (see the text).
3. It is a special blood flow which is established according to the principle of accelerated detoxifying.
4. It is a special threshold value of blood flow rate which exceeds the upper limit of normal blood flow rate in microcirculation of human body.
5. It is the hyperdynamic blood flow which can cause Bernoulli warm shock in clinical practice.
6. It is a theoretical inference which is made according to the conservation law of mass (any inflowing matter is necessary to flow out) and the conservation law of momentum (the velocity is inversely proportional to area) (Feng Yuanzhen, *et al*.,1986)[21], (Professor Emeritus John E et al, 2006)[22].
7. It is the result that the measured results of long-time clinical observation on patients basically consist with the results inferred according to the conservation law of mass and the conservation law of momentum (De Backer et al, 2007) [4].
8. It is classified into three grades of microcirculating hyperdynamic blood flow: suspected (1000-1300 μm/s), highly-suspected (1300-1500 μm/s), and definite (≥1500 μm/s).

In past literatures, the threshold value of hyperdynamic blood flow in sepsis was ever defined. In the literature of [17], the hyperdynamic blood flow in sepsis (V) was defined as blood flow rate of >750 μm/s. Although this threshold value seemed of wide range and even included the blood flow rate of >1000 μm/s, it was defined just for technical reason due to the frame rate of camera and the technical limitation of caecal capillary structure: the frame rate of camera was 30 Hz; the length of vessel bordering crypts was ≤50 μm; only erythrocyte with velocity of ≤750 μm/s could be assessed quantitatively, and the blood flow with velocity of >750 μm/s was denoted as hyperdynamic. Obviously, this definition is not based on results obtained according to any physiological and medical reason; it is essentially different from our definition.

Meanwhile, the literature (A. M. Dondorp et al, 2008) [17] thinks that a hyperdynamic blood flow can not cause oxygen exchange disorder: hyperdynamic blood flow in sepsis patients can not explain tissue anoxia, because oxygen exchange is not influenced even at high erythrocyte velocity and the unloading oxygen is not compromised. This is completely contrary to our explanation and definition of hyperdynamic blood flow according to Bernoulli principle.

## Attachment III

Inclusion criteria for hyperdynamic blood flow sample of clinical sepsis, suggestions/precautions for sampling method and limitations of common space-time method in measurement of hyperdynamic blood flow.

The literature (Paul WG Elbers et al, 2006)[6], (Vanina S et al, 2015)[7], (Vanina S et al, 2015)[8],(Vanina S et al, 2012) [9], (VS Kanoore Edul et al, 2015)[19],(Arnaldo Dubin et al, 2020)[23] reflects that observation results of hyperdynamic blood flow are very controversial and even thinks that the hyperdynamic blood flow does not exist. Actually, it is due to a lack of understanding of the nature of high dynamic blood flow,, a deviation is produced in inclusion criteria and collection method for test sample. Therefore, in order to facilitate the better collection of clinical data, we raise the following suggestions:

A. The nature of hyperdynamic blood flow should be first fully understood through the detoxifying principle and Bernoulli principle. On this basis, an attention should be paid to the following key issues on sample size determination and measurement method while collecting clinical data. Otherwise, wrong conclusion may still be made.
B. The occurrence, development and gradual attenuation of microcirculating hyperdynamic blood flow is a continuous process. If this phenomenon is not understood but it is simply thought that a hyperdynamic blood flow will occur once there is a sepsis, a deviation will be produced for sample collection. The hyperdynamic blood flow only occurs in early or middle period of sepsis and gradually disappears in middle-late and late period of sepsis. Therefore, if this phenomenon is not understood but the sample of sublingual microcirculating blood flow in sepsis of different stage is collected and then analyzed statistically without discrimination, a conclusion that the hyperdynamic blood flow does not exist may be made (Vanina S et al, 2015) [8],(VS Kanoore Edul et al, 2015) [19]. Once it is understood that the occurrence, development and gradual attenuation of microcirculating hyperdynamic blood flow is a continuous process, the heterogeneous phenomenon of microcirculating blood flow often revealed by European scholars can be understood (Paul WG Elbers et al,2006)[6](Can Ince et al, 2018) [10], (Bakker J et al, 2021)[27]. In fact, the heterogeneity is just a fragment at the occurrence, development and gradual attenuation of hyperdynamic blood flow in sepsis. In super-early period of sepsis, the heterogeneous phenomenon can occur only when the hyperdynamic blood flow occurs in only several blood vessels but all of other blood vessels are normal. In early, early-middle and middle period of sepsis, the heterogeneous blood flow will not occur when the hyperdynamic blood flow is completely initiated to fill the whole visual field. In middle-late and late period of sepsis, the hyperdynamic blood flow is gradually attenuated; when there are very many blood vessels with stagnant blood flow, the heterogeneity will occur; at death, blood flow is stagnant in all capillaries, and then the heterogeneity disappears completely. For example: At the International Round-table Conference (2007); three videos of sepsis were reported: In Video 2 (1cc6118-S2) and 3(2cc6118-S3), the heterogeneity did not occur; in Video 4, there was an obvious heterogeneity. In fact, Video 2 (1cc6118-S2) and Video 3 (2cc6118-S3) reflects the condition in early and early-middle period of sepsis that the hyperdynamic blood flow starts the complete initiation; Video 4 (3cc6118-S4) indicates the condition in middle-late period of sepsis that some of hyperdynamic blood flow is attenuated and converted into stagnant blood flow.
C. When cardiac index is >4.0 L/min/m2, hyperdynamic blood flow concurs necessarily not in all cases. As shown by animal experiments of Class I on sheep, they Not fully synchronized, The increase in cardiac index CI and cardiac output CO generally lags behind the occurrence of microcirculatory hyperdynamic blood flow.Therefore, it can not be simply thought that hyperdynamic blood flow will concur necessarily whenever cardiac index is >4.0 L/min/m2 (Paul WG Elbers et al, 2006) [6], (Arnaldo Dubin et al, 2020) [23].
D. In animal experiments, the dose of LPS should be controlled at LPS molding. If its dose is too large, cardiac output will rapidly enter a state of decrease; meanwhile, the hyperdynamic blood flow can not occur in microcirculation. The toxic state of body at this time is a strong stimulus of instant occurrence. At this time, the immune defense mechanism of body does not have time to respond but rapidly enter the state of vasospasm which is caused by overexcitation of sympathetic nerve. However, in sepsis patients, the toxins slowly invade blood; human body can have time to respond and then initiate the immune defense process. At the beginning of our experiments, after LPS injection at too large dose (20 μg/kg in Bama pigs), hyperdynamic blood flow did not occur; after the reduction of LPS dose (5 μg/Kg), hyperdynamic blood flow started to occur. Such phenomenon is also shown in the literature (HENRY M. CRYER,et al,1988)[24].
E. Even though all sample are collected from patients with early sepsis, a deviation is still prone to occur when a blood vessel is selected as measurement sample of hyperdynamic blood flow. In other words, if you do not understand the detoxifying principle of hyperdynamic blood flow, the principle of Xinghuai Feng-Bernoulli warm shock and the threshold value standard of hyperdynamic blood flow, you may not select the blood vessel with highest velocity as measurement sample. Only if the blood flow rate is 1000 μm/s in the blood vessel (For example, vessel A) with highest velocity, oxygen exchange disorder may occur. If this Principle is not understood,Replacing the unique high dynamic blood flow velocity of the vessel A with an average blood flow velocity, the warm shock role of the blood vessel A will be concealed.
F. Advantages and disadvantages of common space-time method: As advantages, this method can automatically measure the velocity; particularly, its measurement is more accurate when the blood flow rate is low. However, when the blood flow rate is >1000 μm/s, very great deviation will be produced. Through the sublingual microcirculation videos of one sepsis patient, we ever compared the measurement results between manual method and space-time method. In manual method, after the macroscopic tracing of leukocyte, the distance was divided by time to obtain mean velocity as 4243 μm/s, which consists with the fluidity results of hyperdynamic blood flow found through macroscopic observation. Through space-time method, mean velocity in the same blood vessel was measured as 0 (Hough transform). As found from spatial-temporal graph, orbital mark direction of erythrocyte had approached to vertical state. According to the principle of spatial-temporal graph, the accuracy of calculated value will be greatly decreased at this time for the following reasons: after the mark direction of spatial-temporal graph approaches to 90°, the projected value of blood cell orbit at time axis will approach to 0 (i.e. tan90°=nonexistent) when the blood flow acceleration is continued; at this time, trigonometric function can not be adopted for accurate calculation of velocity. This is also recorded in the literature (Liuda Wei et al, 2013)[5] (A. M. Dondorp et al, 2008)[17]. The literature(A. M. Dondorp et al, 2008) [17] states as follows: Due to the limitation of image collecting ability, the images can not be captured so rapidly as to quantify the velocity of hyperdynamic blood flow.

At present, manual method is still the most reliable measurement method for hyperdynamic blood flow with velocity of >1000 μm/s. In manual method, the distance of one blood cell is traced visually, and the required time is calculated; even though the next frame is not yet determined as this blood cell, video images can still be played repeatedly to compare and confirm frame by frame; if blood cell is deformed so greatly as not to compare and confirm whether the next frame is still the same cell, another blood cell can be traced and measured instead. As disadvantages, manual method is a semi-automatic operation. However, due to the high reliability, its measured value can be used as reference value of other measurement methods. In addition, manual method can be used to measure the blood flow rate in curved blood vessel.

For many times, the literature (Paul WG Elbers et al, 2006)[6],(Vanina S et al, 2015)[8],(VS Kanoore Edul et al, 2015)[19] has issued the results that hyperdynamic blood flow did not exist, possibly for the following reasons: it may not be understood that only the microcirculating blood flow in patients with early sepsis can be collected; when blood flow rate is >1000 μm/s, measured results through space-time method are very inaccurate. In fact, the literature (Liuda Wei et al, 2013) [5],(A. M. Dondorp et al, 2008) [17] revealed that the space-time method was only applicable to measure the blood flow rate of <1000 μm/s. As shown by our study, initial threshold value of hyperdynamic blood flow was exactly >1000 μm/s. According to the grading criteria, only the blood flow with velocity of >1500 μm/s can be finally determined as hyperdynamic blood flow, which is more beyond the range of measurement accuracy for space-time method.

Above six suggestions aim to prevent the occurrence of large deviation at collection and measurement of hyperdynamic blood flow.

In addition, our definition of hyperdynamic blood flow is applicable for quiescent state (Attachment II). When someone stops after the running for 100 m, the blood flow is definitely accelerated; however, this condition can not be considered as sepsis. The blood flow acceleration at normal exercise can promote the diffusion and exchange of oxygen molecule, because kinetic energy of oxygen molecule is relatively increased to more facilitate its diffusion. However, such situation can occur only under a certain condition: blood flow rate is not too high and can not exceed the upper limit of normal value; there is not great difference between internal and external pressure at capillary. Therefore, the blood flow is accelerated within normal range to more sufficiently implement oxygen exchange, so as to meet the increased demand for oxygen by cells and organs at violent exercise. However, the blood flow acceleration can not be limitless. Once the upper limit of normal value is exceeded, a counteraction will be produced. When the blood flow rate exceeds the upper limit of normal value, The decrease in oxygen partial pressure in true capillaries makes it difficult for oxygen molecules to diffuse out and clinical manifestations of warm shock can occur (i.e. high output and low resistance).

